# Quantitative imaging of loop extruders rebuilding interphase genome architecture after mitosis

**DOI:** 10.1101/2024.05.29.596439

**Authors:** Andreas Brunner, Natalia Rosalia Morero, Wanlu Zhang, M. Julius Hossain, Marko Lampe, Hannah Pflaumer, Aliaksandr Halavatyi, Jan-Michael Peters, Kai S. Beckwith, Jan Ellenberg

## Abstract

How cells establish the interphase genome organization after mitosis is incompletely understood. Using quantitative and super-resolution microscopy, we show that the transition from a Condensin to a Cohesin-based genome organization occurs dynamically over two hours. While a significant fraction of Condensins remains chromatin-bound until early Gl, Cohesin-STAGl and its boundary factor CTCF are rapidly imported into daughter nuclei in telophase, immediately bind chromosomes as individual complexes and are sufficient to build the first interphase TAD structures. By contrast, the more abundant Cohesin-STAG2 accumulates on chromosomes only gradually later in G1, is responsible for compaction inside TAD structures and forms paired complexes upon completed nuclear import. 0ur quantitative time-resolved mapping of mitotic and interphase loop extruders in single cells reveals that the nested loop architecture formed by sequential action of two Condensins in mitosis is seamlessly replaced by a less compact, but conceptually similar hierarchically nested loop architecture driven by sequential action of two Cohesins.

## Introduction

DNA loop extrusion by SMC complexes (structural maintenance of chromosomes) has emerged as a key principle in the spatial organization of chromosomes during interphase and mitosis (Vatskevich *et* al., 2019; Davidson & Peters, 2021). In mitosis, the two pentameric ring-like Condensin complexes I & II, consisting of two shared coiled-coil subunits (SMC2 and SMC4) and three isoform-specific subunits (the kleisin CAP-H or CAP-H2 and two HAWK proteins CAP-D2/3 and CAP-G/2, Hirano & Mitchison, 1994; Hirano et al., 1997) have been shown to be capable of processive DNA loop extrusion (Ganji et al., 2018). Both Condensin I, activated through mitotic phosphorylation and KIF4A (Kimura et al., 1998; Bazile et al., 2010; Tane et al., 2022; Cutts et al., 2024 *Preprint*), and Condensin II, deactivated during interphase by MCPH1 (Houlard et al., 2021) and associating with chromosomes through Ml8BPl in mitosis (Borsellini et al., 2024 *Preprint*), localize to the longitudinal axis of mitotic chromosomes (0no et al., 2003; Hirota et al., 2004). Condensin I & II impact the shape of mitotic chromosomes distinctly, with Condensin II compacting chromosomes axially from prophase onward, and Condensin I compacting chromosomes laterally once it gains access to DNA during prometaphase (0no et al., 2003, 2004; Hirota et al., 2004; Shintomi & Hirano, 20ll; Green et al., 2012). Through their sequential action, the Condensins shape mitotic chromosomes into rod-shaped entities and provide mechanical rigidity (Houlard et al., 20lS) to ensure the faithful segregation of sister chromatids by spindle forces. Based on quantitative and super-resolution imaging, as well as HiC and polymer modelling, it has recently been proposed that Condensins organize mitotic chromosomes into nested loops, with the less abundant and stably binding Condensin II extruding big DNA loops (∼450 kb) already during prophase that are subsequently nested into smaller sub-loops (∼90 kb) by the more abundant and more dynamically associating Condensin I complex after nuclear envelope breakdown (Walther et al., 2018; Gibcus et al., 2018). These Condensin-driven loops are randomly generated across the linear chromosomal DNA molecules, thereby erasing sequence specific interphase structures (Naumova et al., 2013).

In interphase, the two closely related Cohesin complexes Cohesin-STAG1 and Cohesin-STAG2 govern the loop extruder-based genome organization (Wutz et al., 2017, 2020). Like the Condensins, the Cohesins are ring-like protein complexes consisting of two shared coiled-coil subunits (SMC1 and SMC3), a shared kleisin subunit (RAD21, also called SCC1) and one isoform-specific HEAT-repeat subunit (STAG1 or STAG2, Losada et al., 1998, 2000; Sumara et al., 2000). In the presence of the accessory HEAT repeat protein NIPBL, Cohesin complexes can extrude DNA loops (Kim et al., 2019; Davidson et al., 2019) until they are being stalled by the protein CTCF binding to the conserved essential surface of STAG1/2 (Li et al., 2020). CTCF is a zink-finger containing protein that is enriched at its asymmetric cognate binding sites in the genome (de Wit et al., 20lS; Guo et al., 20lS), yielding most efficient stalling of loop extruding Cohesin when arranged in a convergent orientation (Rao et al., 2014). The protein WAPL functions as an un-loader of Cohesin on chromatin, restricting its maximal residence time on chromatin and thereby achieving a constant turnover of DNA loops (Kueng et al., 2006; Wutz et al., 2017). The combined action of these proteins leads to the continuous and dynamic generation of sequence specifically positioned DNA loops in the genome (Rao et al., 2014; Sanborn et al., 20lS; Fudenberg et al., 2016; Gabriele et al., 2022; Mach et al, 2022; Beckwith et al., 2023 *Preprint*), thereby creating more compact domains in the genome termed topologically associated domains (TADs, Nora et al., 2012; Dixon et al., 2012). While their functional role is still an active area of research, TADs have been implicated in the regulation of gene expression through active regulation of enhancer promoter contact frequency (Lupianez et al., 20lS; Zuin et al., 2022).

Similar to the two Condensin isoforms, the Cohesin isoforms STAG1/2 display different expression levels and chromatin residence times, with Cohesin-STAG1 being the less abundant subunit with a long residence time, and Cohesin-STAG2 making up 75% of the total Cohesin pool and being more dynamically bound to chromatin (Losada et al., 2000; Holzmann et al., 2019; Wutz et al., 2020). While the two isoforms share a large portion of common binding sites in the genome and display a certain functional redundancy in the generation of DNA loops, Cohesin-STAG1/2 also have unique binding sites, with Cohesin-STAG1 being preferentially enriched at CTCF binding sites and TAD boundaries, and Cohesin-STAG2 being enriched at non-CTCF sites (Kojic et al., 2018).

While the bona-fide interphase organization and the formation of mitotic chromosomes have been subject to thorough investigation, much less is known about how the interphase organization is rebuilt after mitosis. Previously, the genome-wide reorganization of chromatin has been studied using a combination of HiC and ChiP-seq in cell populations fixed after pharmacological synchronization in a long mitotic arrest. This revealed a slow and gradual transition of the mitotic to the interphase fold over the course of several hours, via an apparently unstructured folding intermediate during telophase that is devoid of Condensin and Cohesin loop extruders, as well as a gradual build-up of TAD structures over the course of several hours during G1 (Abramo et al., 2019; Zhang et al., 2019).

Here, we set out to systematically quantify and map the actions of the Condensin and Cohesin loop extrusion machinery during mitotic exit in single living cells, aiming to characterize the dynamic molecular processes underlying the reformation of the loop-extrusion governed interphase genome organization after mitosis. We find that the switch from mitotic to interphase organization takes about 2 hours in unsynchronized cells, passing a transition state during telophase during which a minimal set of 3 Condensins and Cohesins each are simultaneously bound per megabase of genomic DNA. We find that Cohesin-STAG1 is rapidly imported into the newly formed daughter nuclei alongside CTCF, capable of the formation of large, TAD-scale, loops early after mitosis as a monomer. We find that Cohesin-STAG2 likely also extrudes DNA loops as a monomer, but that it undergoes a concentration-dependent dimerization on chromatin upon its full import into the nucleus. Based on our quantitative imaging data, we can infer that this phenomenon is a result of the high occupancy of 8 chromatin-bound Cohesin-STAG2 per megabase in late G1, leading to frequent encounters of neighboring complexes that lead to a nested/stacked arrangement of extruded loops. Surprisingly, we also find that CTCF is increasingly stabilized on chromatin throughout G1 due to its increasing interaction with the two Cohesin complexes. Based on these data, we propose a double-hierarchical loop model to generate interphase genome architecture after mitosis, in which the two interphase Cohesin loop extruders sequentially build a nested arrangement of large and then small DNA loops, conceptually similar to how Condensins have been suggested to drive mitotic genomic organization.

## Results

### The transition from mitotic to interphase loop extruders occurs over two hours after mitosis and requires nuclear import

To examine the time required to complete the switch from mitotic to interphase loop extruder genome organization (Fig. 1A), we made use of human HeLa Kyoto (HK) homozygous knock-in cell lines in which all alleles of the endogenous genes for the kleisin subunits of Condensin I (NCAPH), Condensin II (NCAPH2) and the HEAT-repeat subunits of Cohesin-STAGl (STAGl) and Cohesin-STAG2 (STAG2) have been tagged with GFP (Walther et al., 2018; Cai et al., 2018). After a single S-phase synchronization, we performed continuous FCS-calibrated 4D live-cell imaging (Politi et al., 2018, Fig. 1B, Suppl. Fig. 1A) through two subsequent cell divisions with 10-minute time-resolution, using SiR-Hoechst and extracellular Dextran to label nuclear and cell volumes, respectively (Fig. 1C). Computational 3D segmentation of these cellular landmarks (Cai et al., 2018, Suppl. Fig. 1B), combined with automatic cell tracking allowed us to align single cell trajectories from one anaphase to the next, and calculate absolute protein concentrations and copy numbers throughout a full cell cycle (Fig. 1D, Suppl. Fig. lC-G).

**Figure 1.**
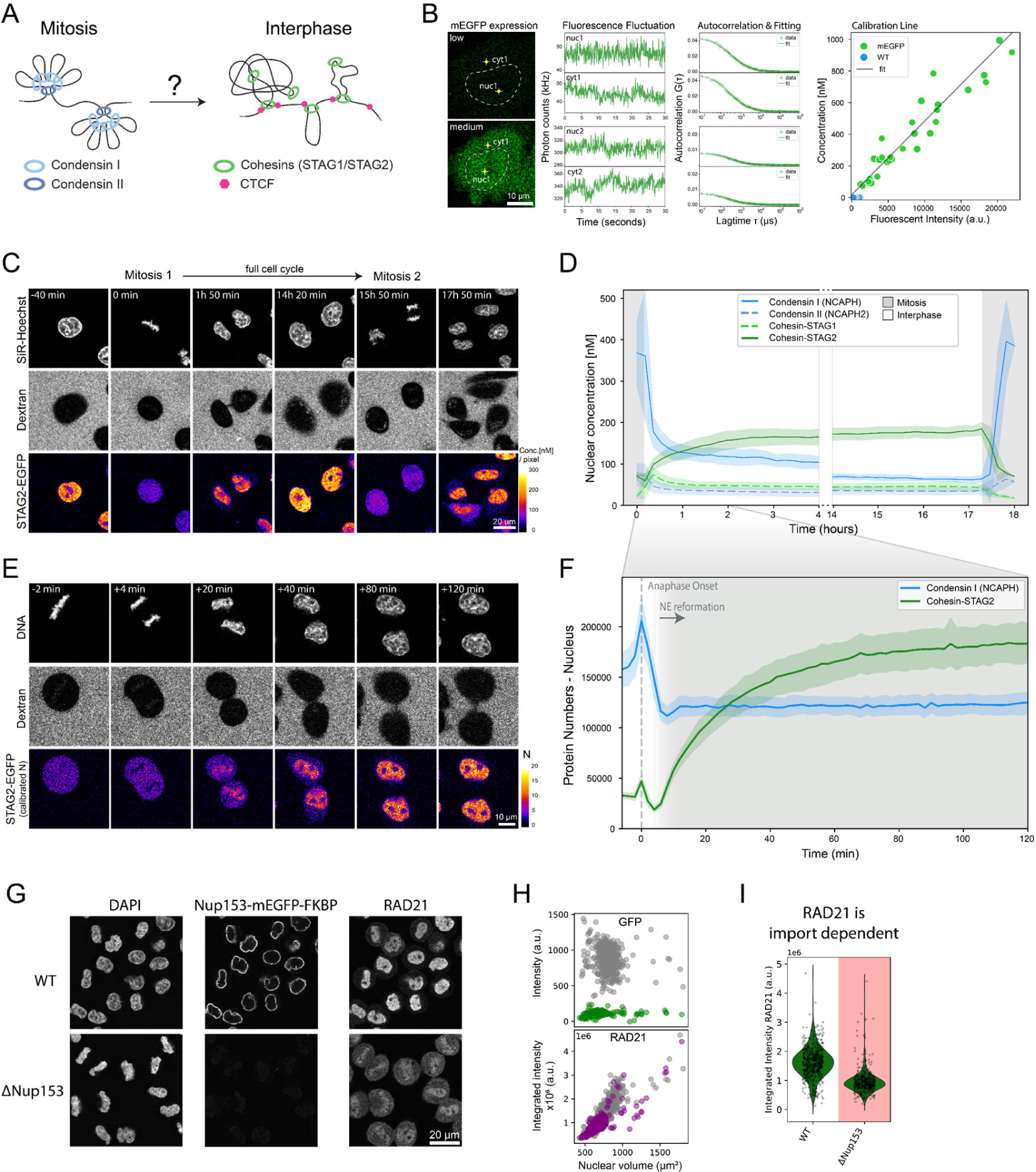
FCS-calibrated imaging of Cohesin isoforms shows that re-organization of loop extrusion during mitotic exit takes about 2h after anaphase onset. **A)** Schematic of current loop-extrusion based models of mitotic and interphase genome organization. Condensin I and II build nested mitotic loops. Cohesins build topologically associating domains delimited by the boundary factor CTCF during interphase. **B)** Fluorescence intensity calibration using fluorescence correlation spectroscopy (FCS). Fluorescence intensity and photon count fluctuation measurements are performed in cells expressing varying amounts of monomeric EGFP. An autocorrelation function simulating particle diffusion through the effective detection volume is fit to the autocorrelated photon count signal, enabling estimation of protein number in the effective detection volume and therefore the calibration of fluorescent intensities to absolute protein concentrations (Politi et al., 20l8). **C)** Imaging of genome-edited HK cells with homozygously EGFP-tagged Cohesin-STAG2 throughout 2 consecutive mitoses. Fluorescent intensities (FI) measured in the second mitosis were set to the average protein concentration (C[nM]) during metaphase as measured by FCS-calibrated imaging of the same cell line during mitotic exit imaging (see E)). Protein concentrations of all other timepoints were adjusted based on relative differences of measured fluorescence intensities. **D)** Nuclear concentration throughout an entire cell cycle ranging from one anaphase to the next displayed for genome-edited HK cells with homozygously (m)EGFP-tagged Cohesin-STAGl (n = 20 cells) and Cohesin-STAG2 (n = l3 cells), as well as Condensin I (NCPAH, n = 8 cells) and Condensin II (NCAPH2, n = 2l cells). Inset shows focus of imaging on first two hours after mitosis performed with higher temporal resolution. Error bands represent 9S% confidence interval. **E)** FCS-calibrated imaging of genome-edited HK cells with homozygously EGFP-tagged Cohesin-STAG2 throughout mitotic exit. A total of 7S 3D stacks with 2-minute intervals is triggered after successful automated identification of metaphase cells based on SiR-Hoechst staining. Fluorescence intensity calibration by FCS allows for the conversion of measured fluorescent intensities (FI) to absolute protein concentrations and protein numbers (N) per unit volume. **F)** Absolute protein numbers co-localizing with chromatin/the two daughter nuclei displayed for genome-edited HK cells with homozygously (m)EGFP-tagged Cohesin-STAG2 (n = ll cells) and Condensin I (NCPAH, n = l4 cells). Reformation and full establishment of the nuclear envelope as determined by Lamin B receptor (Suppl. Fig. lN) is indicated through grey background. Error bands represent 9S% confidence interval. **G)** Fluorescent micrographs of early Gl cells (∼4S min past mitosis) stained with DAPI in WT or ΔNuplS3 condition. **H)** Average fluorescent intensity plots per 3D-segmented nucleus in grey (WT) or colored (ΔNuplS3). ΔNuplS3 nuclei do not expand in size, show no residual NuplS3 intensity and show a clear reduction in RAD2l intensity inside the nuclear lumen. **I)** Average fluorescent intensity of early Gl cells in WT or ΔNuplS3 condition stained for RAD2l. S0% drop in mean RAD2l intensity after 7S min release time. 33-48% reduction in average fluorescent intensity after 4S min release time (not shown). Changes above/below 20% are considered a significant change.

As expected, both Condensin isoforms were concentrated on mitotic chromosomes and Condensin II maintained a stable nuclear concentration after division (Fig. 1D). Surprisingly, we found that while the high Condensin I concentration of 380 nM on chromosomes dropped sharply after segregation, it did not become completely cytoplasmic but maintained a concentration of lS0 nM in the two newly formed interphase nuclei, where it then became diluted slowly with nuclear growth (Fig. 1D). Photobleaching of this nuclear Condensin I pool in interphase revealed that it moves freely in the nucleus and does not exchange with the cytosolic pool (Suppl. Fig. 1H). Quantitative full cell cycle imaging showed that the nuclear pools of NCAPH/2 could in principle form complete Condensin complexes with the shared Condensin subunit SMC4, which is present inside the nucleus in sufficient numbers (Suppl. Fig. 1E).

Conversely to the sharp reduction in chromosomal Condensin I, the two Cohesin isoforms STAG1/2 that are key for interphase genome organization became enriched inside the nucleus after anaphase and reached essentially constant nuclear concentrations throughout the entire interphase (Fig. 1D). This was also found to be true for CTCF (Suppl. Fig. 1E), revealing homeostatic stable nuclear concentrations of these factors and no doubling of interphase loop extruders with DNA replication, consistent with previous reports that showed uncoupling of nuclear growth from DNA replication (0tsuka et al., 2016). We found that the Cohesin isoform STAG1 only makes up 25% of the total Cohesin pool, consistent with previous studies (Fig. 1D, Losada et al., 2000; Holzmann et al., 2019). Interestingly, Cohesin-STAG1 displayed rapid and complete nuclear localization shortly after mitosis, followed by an equilibration of its nuclear concentration upon nuclear expansion (Fig. 1D, Suppl. Fig. 1D). The 3-fold more abundant Cohesin-STAG2, however, reached stable nuclear concentrations only about two hours after mitosis (Fig. 1D, Suppl. Fig. 1I-N).

Having characterized the chromosomal/nuclear concentration changes of the four loop extruders and CTCF throughout the cell cycle, we next focused our analysis on the transition between Condensin and Cohesin occupancy on the genome during the first 2 hours after mitosis, increasing the time-resolution of our FCS-calibrated 4D imaging to 2 minutes (Fig. 1E). This detailed kinetic analysis revealed that the number of Condensin I proteins associating with chromatin rapidly dropped after its peak during anaphase (Fig. 1F). This drop, however, ceased at the time of reformation of the nuclear envelope, 6 minutes after A0 (Suppl. Fig l0), that creates a permeability barrier and apparently retains the remaining Condensin I molecules inside the newly formed nucleus. Cohesin-STAG2 started its nuclear enrichment precisely from the time of nuclear envelope assembly, and required 2 hours until complete nuclear enrichment (Fig. 1F, Suppl. Fig. 1N). To test if the Cohesin accumulation required nuclear import, we acutely degraded degron knocked-in NuplS3, an essential component of nuclear pore complexes (Fig. 1G, Suppl. Fig. 1Q-S). We found that both Cohesin and CTCF levels inside the nucleus were significantly reduced in NuplS3 depleted cells early after mitosis (Fig. 1G-I, Suppl. Fig. 1T), showing that both factors require functional nuclear pores to reach the genome.

### Condensins and Cohesins bind simultaneously, yet independently, to the early Gl genome at 3 complexes per megabase DNA

Given that a significant number of both Condensin complexes are still present inside the newly formed daughter cell nuclei when Cohesins start to be imported (Fig. lF, Table l), we wanted to go beyond nuclear concentration and protein numbers and ask how much of the mitotic and interphase loop extruding complexes are bound to chromatin after mitosis and could thus be actively engaged in extrusion. To quantify binding, we used fluorescence recovery after photobleaching (FRAP, Suppl. Fig. 2A) of Condensin I and II on the metaphase plate and in the newly formed nucleus. Half nuclear photobleaching indicated that a significant fraction of Condensins remains chromatin-bound in early G1 (Suppl. Fig. 2 B-D), while at the same time a large fraction of the newly imported Cohesins are already bound (Suppl. Fig. 2F). To assay changes in the chromatin-bound fraction of Condensins and Cohesins quantitatively and in a highly time-resolved manner during mitotic exit, we used a rapid spot-bleach assay monitoring fluorescence depletion from a femtoliter-sized chromatin volume during 30 second continuous illumination with a diffraction limited focused laser beam (Fig. 2A, Suppl. Fig. 2G-I). In this assay, the chromatin-bound protein fraction is bleached, while the unbound fraction recovers from the excess soluble nuclear pool outside the small bleach spot. This approach thus provides a rapid measure for the bound fraction of GFP-tagged proteins on chromatin that can be carried out repeatedly in a single living cell without interfering with mitotic progression (see Supplementary Table 1&2 for more detail and comparison to classical FRAP).

**Figure 2.**
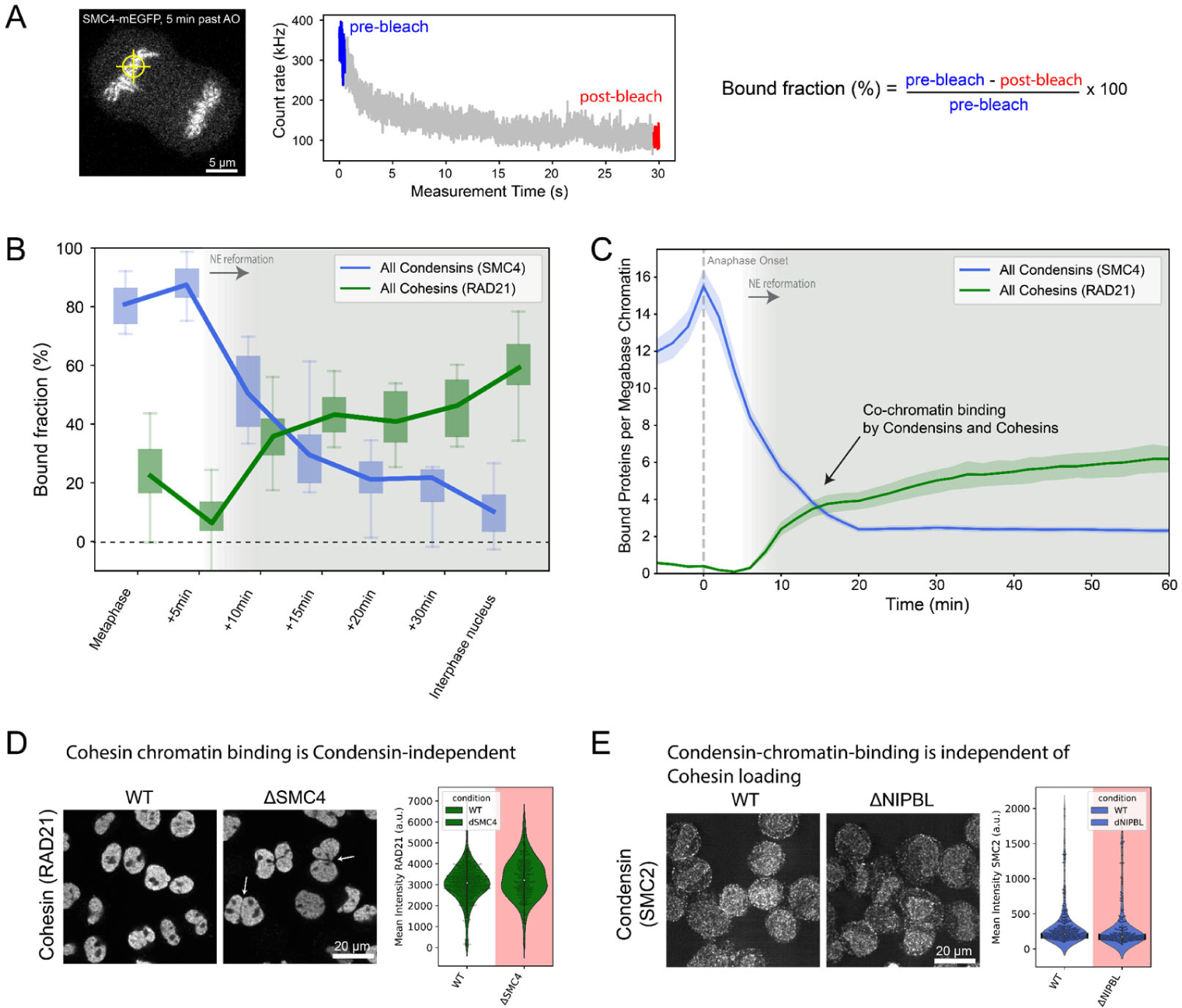
Condensins and Cohesins co-occupy chromatin during telophase and early G1, as revealed by time-resolved bleaching. **A)** Illustration of the spot-bleach assay. Genome edited HK cells homozygously expressing (m)EGFP tagged Condensin and Cohesin subunits are illuminated at a single spot on chromatin for a total duration of 30 seconds and the resulting fluorescent intensity is continuously measured. The chromatin-bound fraction of a given protein of interest is calculated based on the mean fluorescent intensity of the first and last S00 msec. Exemplary image and bleach data is shown for the common Condensin subunit SMC4. **B)** The fraction of chromatin-bound Condensins (SMC4) and Cohesins (RAD2l) determined using the spot-bleach assay at different timepoints during mitotic exit. Every bar plot represents at least l0 individual datapoints measured in l0 separate cells. **C)** Absolute number of proteins bound to chromatin were determined by multiplication of chromatin bound fractions shown in B with absolute protein numbers co-localizing with chromatin (n(SMC4) = 2l cells, n(RAD2l) = l8 cells) as determined in Fig. lE&F and displayed as per-megabase-count assuming an equal distribution of the proteins on the entire 7.9 Mb HeLa genome (Landry et al., 20l3). Grey background indicates reformation of nuclear envelope. Error bands represent 9S% confidence interval. **D)** Fluorescent micrographs and quantification of early Gl cells in WT condition or after degradation of the isoform-shared Condensin subunit SMC4. Cells were pre-extracted for l minute prior to fixation and were stained for RAD2l. SMC4 depletion caused a delay in cell division as well as major cell division errors (see merged daughter nuclei in fluorescent micrograph indicated by arrow). Time of release from Nocodazole block had to be increased to 60-70 minutes to fix cells in early Gl stage. Difference in mean fluorescence intensity: 8-l2.S%. Changes above/below 20% are considered a significant change. **E)** Fluorescent micrographs and quantification of early Gl cells in WT condition or after degradation of the Cohesin loader NIPBL. Cells were pre-extracted for l minute before fixation and were stained for SMC2. Difference in mean fluorescence intensity: ∼lS%. Changes above/below 20% are considered a significant change.

**Table 1.**
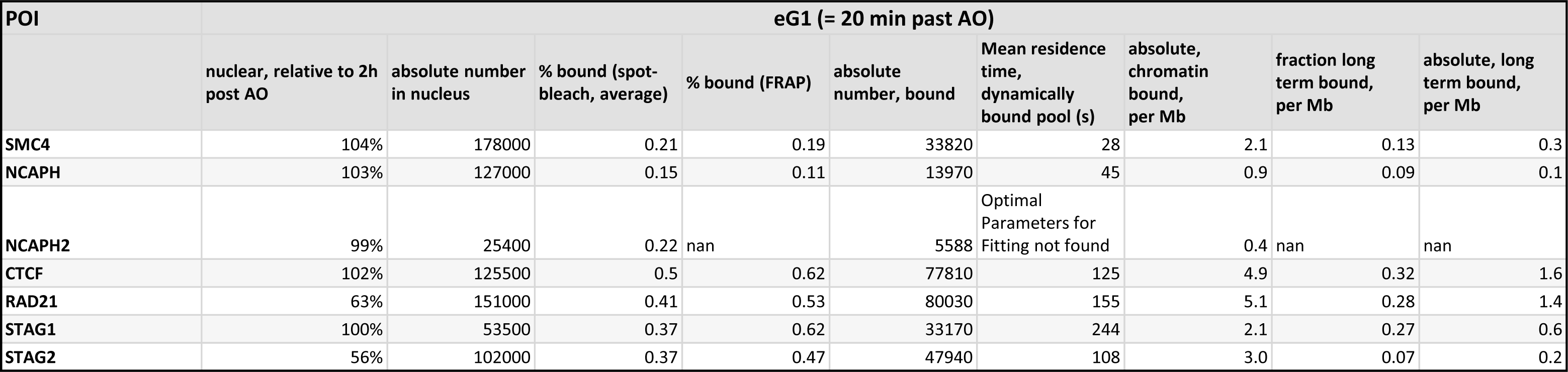
Quantitative data summary: early G1.

| POI | eG1 (= 20 min past AO) |  |  |  |  |  |  |  |  |
| --- | --- | --- | --- | --- | --- | --- | --- | --- | --- |
|  | nuclear, relative to 2h post AO | absolute number in nucleus | % bound (spot-bleach, average) | % bound (FRAP) | absolute number, bound | Mean residence time, dynamically bound pool (s) | absolute, chromatin bound, per Mb | fraction long term bound, per Mb | absolute, long term bound, per Mb |
| SMC4 | 104% | 178000 | 0.21 | 0.19 | 33820 | 28 | 2.1 | 0.13 | 0.3 |
| NCAPH | 103% | 127000 | 0.15 | 0.11 | 13970 | 45 | 0.9 | 0.09 | 0.1 |
| NCAPH2 | 99% | 25400 | 0.22 | nan | 5588 | Optimal Parameters for Fitting not found | 0.4 | nan | nan |
| CTCF | 102% | 125500 | 0.5 | 0.62 | 77810 | 125 | 4.9 | 0.32 | 1.6 |
| RAD21 | 63% | 151000 | 0.41 | 0.53 | 80030 | 155 | 5.1 | 0.28 | 1.4 |
| STAG1 | 100% | 53500 | 0.37 | 0.62 | 33170 | 244 | 2.1 | 0.27 | 0.6 |
| STAG2 | 56% | 102000 | 0.37 | 0.47 | 47940 | 108 | 3.0 | 0.07 | 0.2 |

We then used this assay to monitor changes in chromatin-binding of Condensin and Cohesin every 5 minutes after exit from mitosis. We found that while all Condensins (using the isoform-shared subunit SMC4-mEGFP) progressively dissociated from chromatin during telophase and early G1, they retained a significant chromatin bound fraction of around 2S% lS minutes after A0 (Fig. 2B). This reduction in bound fraction, also following nuclear envelope reformation, was consistent for both Condensin isoforms, as shown by time-resolved spot bleaching using isoform-specific NCAPH and NCAPH2 subunits (Suppl. Fig. 2K). By contrast, we found that the fraction of bound Cohesins (using isoform shared subunit RAD21-EGFP) increases continuously following nuclear envelope reformation (Fig. 2B), reaching a bound fraction of about 40% lS min after A0. Again, this increase in binding was consistent for both Cohesin isoforms (using isoform specific STAG1/2 subunits (Suppl. Fig. 2K).

0ur quantitative real time analysis of chromatin binding in single dividing cells provides clear evidence for co-occupancy of chromatin by Condensin and Cohesin complexes throughout telophase and early G1. Combining the bound fraction measurements by FRAP with the protein numbers measured by FCS-calibrated imaging (e.g. Fig. 1F, Suppl. Fig. 1P) allows us to calculate the number of proteins bound to genomic DNA (Fig. 2C, Suppl. Fig. 2L). This analysis shows that in early Gl, lS minutes after A0, the same number of around three Condensin and Cohesin complexes are simultaneously bound per megabase of genomic DNA (Fig. 2C). Could this simultaneous binding of mitotic and interphase loop extruders be functionally interlinked? To test this, we probed if the chromatin localization of Condensins and Cohesins in early G1 depends on each other’s presence, using AID-degron knock-in cell lines for the isoform-shared Condensin subunit SMC4 (Schneider et al., 2022) and the Cohesin-chromatin-loader NIPBL (Mitter et al., 2020). In these cells, we could acutely degrade the degron tagged proteins during mitosis (Suppl. Fig. 2J), and ask if they are required for the other complex to associate with chromatin by subsequent immunofluorescence staining for the non-degraded Condensin or Cohesin complex. This analysis did not show major differences in chromatin association of Condensin after NIPBL or Cohesin after SMC4 depletion, respectively (Fig. 2D&E). This suggested that while mitotic and interphase loop extruders bind to chromatin simultaneously in very similar numbers during G1, they do so independently.

### Cohesin-STAGl and CTCF are simultaneously imported immediately after mitosis and sufficient to build the first interphase hallmarks in genome structure

0ur full cell cycle data showed clear differences in the time required for complete nuclear import of the two Cohesin isoforms, with STAG1 reaching maximal nuclear concentration within only 10 mins, while STAG2 reached steady state only after over two hours (Fig. 1D, Suppl. Fig. 1N). To get a first insight into which complex might functionally be more important for early G1 genome architecture, we compared these kinetics with the boundary factor CTCF using an endogenous CTCF-EGFP knock-in cell line (Cai et al., 2018). Calibrated full cell cycle imaging showed a strikingly similar kinetic signature of its nuclear concentration changes compared to Cohesin-STAG1, reaching an approximately 2.5 times higher steady state concentration in interphase (Suppl. Fig lE). We therefore compared the nuclear import kinetics of Cohesin-STAG1 and CTCF relative to the slower accumulating Cohesin-STAG2 with high time-resolution after mitotic exit, using our FCS-calibrated 4D imaging setup. Strikingly, we found that CTCF displayed indistinguishable import kinetics as Cohesin-STAG1 while Cohesin-STAG2 was imported at a much lower rate (Fig. 3A, Suppl. Fig. 3A).

**Figure 3.**
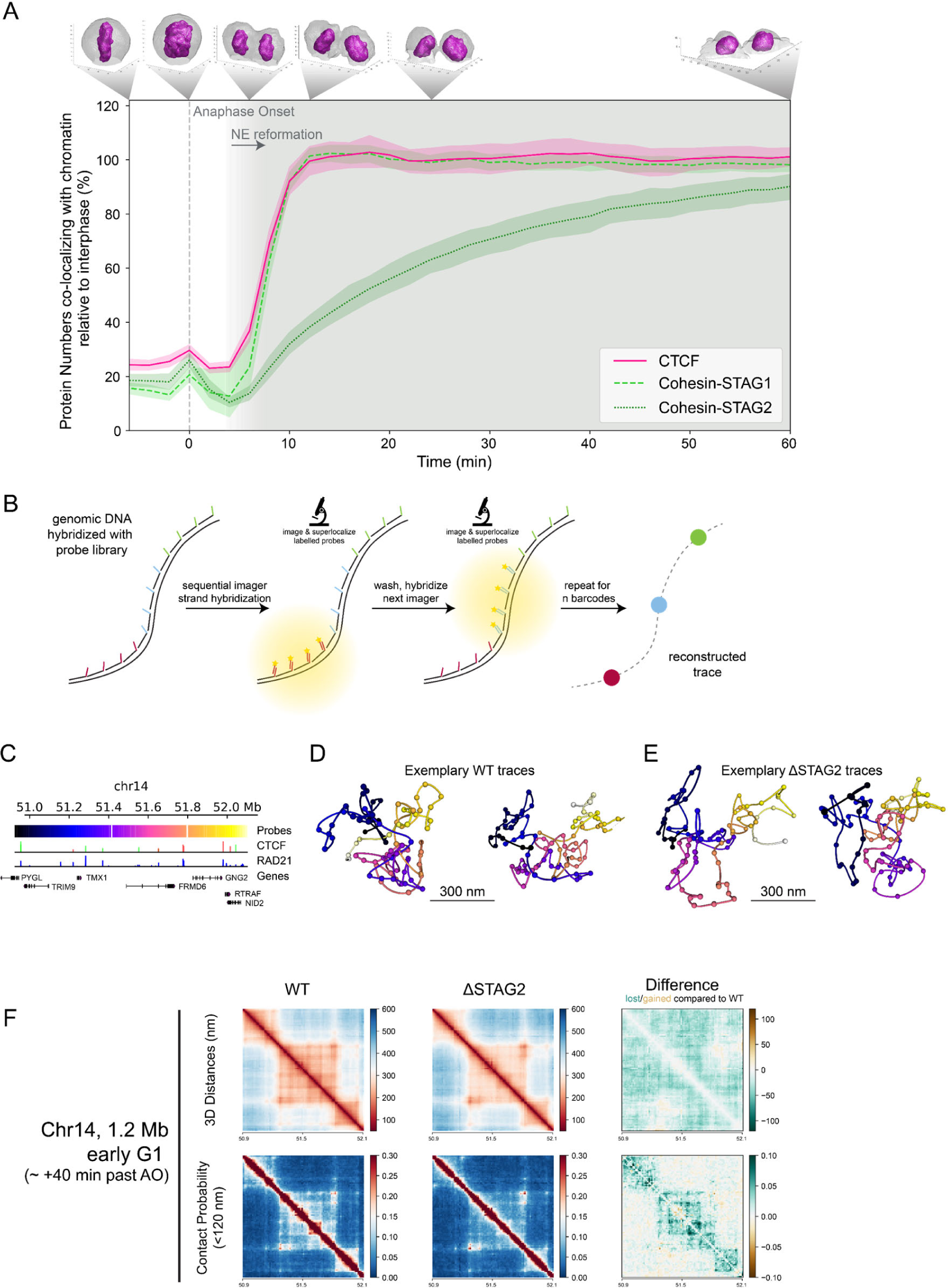
Cohesin-STAG1 and CTCF cooperate to form interphase TAD structures after mitosis. **A)** FCS-calibrated protein numbers co-localizing with chromatin displayed for genome-edited HK cells with homozygously EGFP-tagged Cohesin-STAGl (n = 2S cells), Cohesin-STAG2 (n = ll cells) and CTCF (n = lS cells) relative to the measurement 2 hours after anaphase onset. Error bands represent 9S% confidence interval. **B)** Scheme explaining LoopTrace chromatin tracing workflow. Fixed cells were subjected to single strand resection via exonuclease treatment (RASER) for maximal structure-preservation and subsequent hybridization with a tiled FISH library. Every FISH-probe contains a non-genome-complementary docking handle that can be hybridized with a fluorescently labelled imager strand to read out the 3D location of a genomic locus (Beckwith et al., 2023 *Preprint*). **C)** 0verview of the traced l.2 megabase locus on chromosome l4 with genes as well as ChIP-seq binding sites for RAD2l and CTCF (from the ENC0DE portal (Sloan et al., 20l6, https://www.encodeproject.org/) with the following identifiers: ENCFF239FB0 (RAD2l), ENCFFlllRWV (CTCF); CTCF directionality annotations from Rao et al., 20l4). **D-E)** Exemplary chromatin traces of WT (E) or ΔSTAG2 (F) early Gl cells. **F)** Distance and contact matrices of a l.2 megabase region on chromosome l4 locus traced at a genomic resolution of l2 kb in early Gl cells with and without Cohesin-STAG2. Differences between WT and ΔSTAG2 are highlighted for distance and contact probability maps.

The simultaneous import of Cohesin STAG1 and CTCF is consistent with a functional interaction on chromatin immediately after nuclear reformation. To test if the two proteins are bound to chromatin, we performed real time spot-bleach, as well as FRAP measurements of CTCF, to compare its binding to chromatin with Cohesin STAG1 early after mitotic exit (Suppl. Fig. 2E&L). This analysis revealed that when Cohesin-STAG1 and CTCF reach their maximum concentration, about 2 Cohesin-STAGl and S CTCF molecules are bound per megabase of genomic DNA (Table 1). Two actively extruding Cohesin-STAGl complexes per megabase of genomic DNA would in principle explain the frequency of compact topologically associated domain (TAD) structures that have been estimated at l.S TADs/Mb using biochemical approaches previously (Wutz et al., 2020). To test directly, if Cohesin STAG1 without Cohesin STAG2 is indeed sufficient to create the first more compactly folded G1 genome structures in single cells, we took advantage of our recently developed nanoscale DNA tracing method LoopTrace, enabling us to inspect individual 3D DNA folds as well as ensemble averages with precise physical distance measures (Beckwith et al., 2023 *Preprint*, Fig. 3B, Suppl. Fig. 3B). We traced three independent 1.2 megabase long genomic regions predicted to contain TADs, in 3D at 12 kb genomic and 20 nm spatial resolution (Fig. 3C, Suppl. Fig. 3E&F). 0ur single cell DNA traces could indeed readily identify compact 3D DNA folds already in single early G1 cells (Fig. 3D). Depletion of Cohesin-STAG2 during the prior mitosis (Suppl. Fig. 3B&C) did not influence the overall genomic size of these domains, but led to some reduction in internal loop nesting and slight physical decompaction (Fig. 3E, Suppl. Fig. 3D), which was also clear when comparing pairwise physical 3D distance maps of these regions from hundreds of control or STAG2 depleted cells (Fig. 3F, Suppl. Fig. 3G&H). We conclude that Cohesin-STAG1 and CTCF are imported with identical kinetics rapidly after mitosis and are sufficient to build the first compact looped interphase structures in single G1 cells, equivalent to biochemically detected TADs in cell populations.

### Cohesin-STAGl and CTCF become increasingly stably bound to the genome throughout Gl

To investigate the interplay of Cohesin-STAG1 and Cohesin-STAG2 at later times after mitosis, we performed FRAP measurements during G1 (2-Sh past A0) and compared them to our measurements shortly after mitosis (Fig. 4A). We found that the chromatin-bound fraction for both Cohesin isoforms as well as CTCF significantly increased in later G1 (Suppl. Fig. 4A). A single exponential function with an immobile fraction fit the fluorescence equilibration kinetics of all proteins well (Fig. 4B) and allowed us to determine the dynamically chromatin-bound protein fraction, its residence time, as well as the stably bound fraction that did not exchange dynamically during our measurement time (Suppl. Fig. 4B&C, Tables 1&2). While the average residence time of the dynamically bound pool of Cohesin isoforms (STAG1: 4 min, STAG2: 2 min and CTCF: 2 min) remained unchanged from early to late G1 (Suppl. Fig. 4B), we measured a significant increase in the stably chromatin-bound fraction for Cohesin-STAG1 and CTCF, reaching up 30-40% of the total protein (Fig. 4C&E, Suppl. Fig. 4C). Cohesin-STAG2 also displayed a significant increase in its stably chromatin-bound fraction, however reaching less than 10% of the total protein pool (Fig. 4D). While it has been previously reported that Cohesin-STAGl chromatin binding can be stabilized by CTCF (Wutz et al., 2020), whether CTCF’s own binding is reciprocally affected by the presence of Cohesin has not been investigated. To test if CTCF’s increasingly stable binding in G1 depends on Cohesin, we acutely depleted the isoform-shared subunit RAD21 (Suppl. Fig. 4D-G), which resulted in a significant reduction of stably chromatin-bound CTCF, which could be rescued by RAD21 overexpression (Fig. 4F). This shows that Cohesin is necessary and sufficient to stabilize CTCF’s interaction with chromatin in Gl.

**Figure 4.**
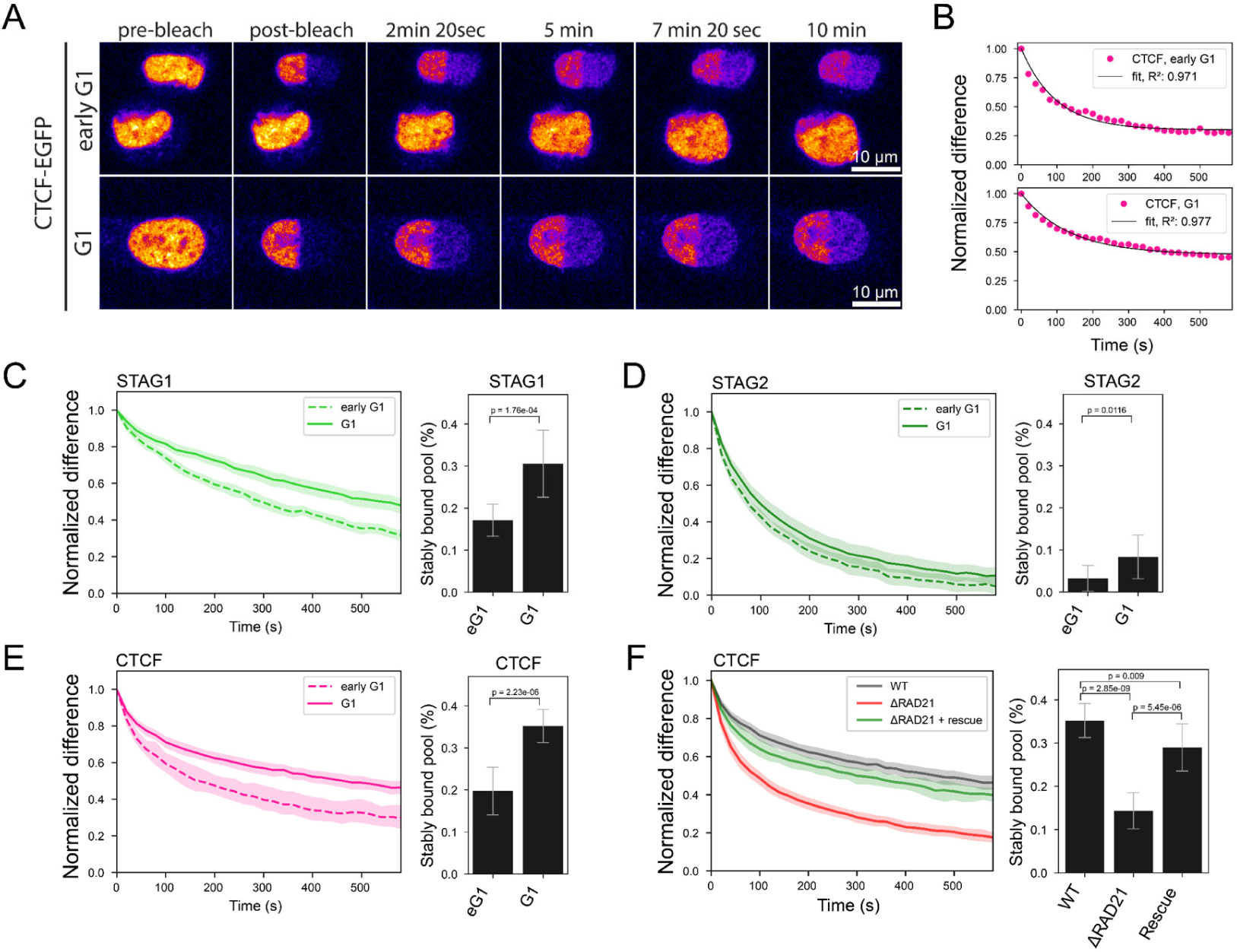
Fluorescence Recovery of Cohesin isoforms and CTCF. **A)** Fluorescence recovery after photobleaching (FRAP) was performed by bleaching half of a nucleus in early Gl cells (20-40 minutes after anaphase onset) or later Gl cells selected by nuclear volume. **B)** FRAP shown for genome-edited HK cells with homozygously EGFP-tagged CTCF. Difference between the bleached an unbleached region is normalized by the maximal difference at time t=0 after bleaching. Black line indicates the data fit by a single-exponential function with immobile fraction. Single exponential functions with immobile fraction also fit the FRAP recovery of RAD2l, STAGl/2 well. **C-E)** FRAP measurements using homozygous EGFP-knock-in HK cell lines in earlyGl and Gl cells, respectively. Bar plots display the fraction of protein that is stably bound to chromatin. Two-sample t-test was used for calculating significance levels. Error bars show standard deviation. **C)** Cohesin-STAGl (early Gl: n = l0 cells, Gl: n = 9 cells) **D)** Cohesin-STAG2. (early Gl: n = l0 cells, Gl: n = l3 cells) **E)** CTCF. (early Gl: n = 9 cells, Gl: n = l0 cells) **F)** FRAP measurements of endogenous CTCF with WT levels of RAD2l, after degradation of endogenous RAD2l, and after rescue of RAD2l degradation by exogenous RAD2l expression for at least 24 hours. Bar plots display the fraction of protein that is stably bound to chromatin. (CTCF WT: n = l0 cells, CTCF dRAD2l: n = 9 cells, CTCF dRAD2l rescue: n = l0 cells). Data from CTCF-EGFP knock-in line is used as WT reference as it displays WT expression levels of RAD2l. The double-knock-in line Rad2l-EGFP-AID CTCF-Halo-3xALFA #C7 displayed leaky degradation of RAD2l, reducing CTCF-chromatin binding already in -IAA cells (see Suppl. Fig. 4G and methods).

**Figure 5.**
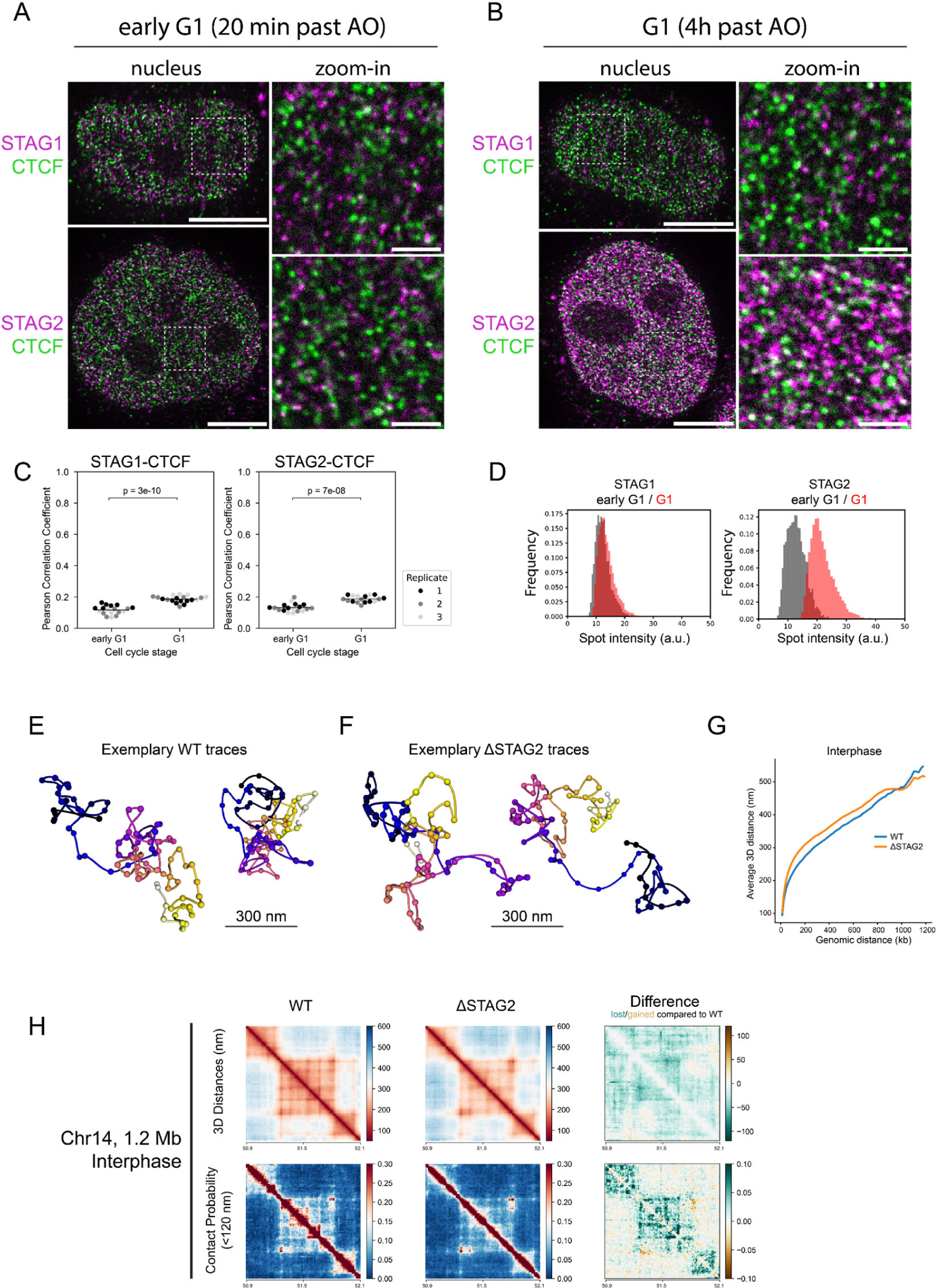
STED super-resolution imaging of Cohesin isoforms and CTCF. **A-B)** Exemplary STED images showing Cohesin-STAG1/2 (magenta) and CTCF (green) in early G1 (A) and G1 (B) nuclei (scalebar: 5 µm) and zoom-ins (scalebar: 1 µm). **C)** Co-localization analysis of Cohesin-STAG1/2 with CTCF using the Pearson Correlation Coefficient of segmented nuclei (n ≥ l7). Differences in STAG-CTCF co-localization in early Gl compared to Gl are significant as assessed by independent two-sample t-test. **D)** Mean intensity of segmented STAGl/2 spots in STED images of replicate 3. Same results are observed in replicate l and 2. Formal significance tests are meaningless due to large sample size. Median intensities of the mean spot intensity distributions are: STAGl eGl: l2.02, STAGl Gl: l3.06, STAG2 eGl: l2.62, STAG2 Gl: 2l.02 (arbitrary intensity units). **E-F)** Exemplary chromatin traces of WT (E) or ΔSTAG2 (F) interphase cells. **G)** Scaling plot of Chrl4 l.2 megabase region sampled at l2 kb resolution in WT or ΔSTAG2 interphase cells. Traces from ΔSTAG2 are on average less compact compared to WT. **H)** Distance and contact matrices of a l.2 megabase region on chromosome l4 locus traced at a genomic resolution of l2 kb in interphase cells with and without Cohesin-STAG2. Differences between WT and ΔSTAG2 are highlighted for distance and contact probability maps. WT data from l, ΔSTAG2 data from 2 independent technical replicates (392 and 6l0 traces, respectively).

### Cohesin-STAG2 completes its nuclear import after 2 hours and exhibits concentration dependent dimerization on the genome in Gl

To test directly whether the observed interdependent increase in stable binding of both CTCF and Cohesins is due to increased complex formation between these proteins on chromatin, we performed STED super-resolution imaging of CTCF and the Cohesin isoform specific subunits STAG1 and STAG2 during early and late G1. To achieve high and comparable labeling efficiency of the different Cohesin isoforms, we used our homozygous knock-in cell lines for STAG1/2-EGFP and detected both with the same GFP-nanobody, while using a specific antibody to detect endogenous CTCF as a reference (Fig. 5A&B, Suppl. Fig. 5A-C). Having calculated the number of chromatin complexes from our combined concentration imaging and FRAP data allowed us to estimate the labeling efficiency of our super-resolution imaging by counting the individual labeled fluorescent spots to about 60-80% for the two Cohesin-isoforms (Suppl. Fig. 5D-F), very similar to our previous labeling efficiencies of this GFP nanobody (Thevathasan et al., 2019). Given that we could resolve the expected number of Cohesin complexes as individual fluorescent spots, the large majority of the labelled STAG1/2 proteins in early G1 therefore most likely represent monomeric Cohesin complexes.

Colocalization of either STAG1 or STAG2 with CTCF resulted in comparable spatial correlations that in both cases increased slightly but significantly from early to late G1, indicating an increase in Cohesin-CTCF complex formation for both isoforms (Fig. 5C). This increased colocalization was not due to the still ongoing accumulation of STAG2 in the nucleus, as shown by image simulations with random protein distributions at realistic densities (Suppl. Fig. 5H&I). 0ur data thus suggests that CTCF associates with both STAG1, that enters the nucleus early, and STAG2, that completes its import later and eventually becomes the more abundant Cohesin isoform. This finding is also in line with the fact that also Cohesin-STAG2 becomes more stably bound to chromatin in late G1 (Fig. 4D).

The ability to detect both Cohesin isoforms at the single complex level with high labeling efficiency in early G1 also put us in a position to use the intensity of Cohesin spots to ask if we can detect multimerization of the Cohesins as the cell cycle progresses, which would be expected with the formation of closely stacked or nested loops. Interestingly, this analysis revealed that while the average spot intensity and total number of spots detected did not change for Cohesin-STAG1 between early and late G1, the STAG2 spot intensity increased about 2-fold between early and late G1 (Fig. 5D, Suppl. Fig. 5G) and correlated with a 2-fold drop in the number of STAG2 spots detected compared to our expectation from quantitative live imaging (Suppl. Fig. 5F). When it has reached its maximum concentration in late G1, Cohesin-STAG2 thus appears to associate with the genome in pairs of molecules that are no longer resolvable individually by STED microscopy, that has a lateral precision of around 60 nm. With an estimated extension of single Cohesin complexes of 50 nm, they must therefore be very closely adjacent to each other or form dimers to result in single STED spots with doubled intensity.

Why might Cohesin-STAG2 form closely adjacent pairs of complexes only at the end of G1? To test if its self-association is concentration dependent, we performed partial depletion of degron tagged Cohesin-STAG2. This indeed shifted the average spot brightness in late G1 back to a value of nanobody monomers (preliminary data, not shown), supporting a concentration rather than for example a cell cycle driven dimerization of Cohesin-STAG2 on DNA. In fact, our quantitative imaging data of the increasing numbers of chromatin-bound Cohesins after mitosis provides a quantitative explanation for the concentration-dependent dimerization of Cohesin-STAG2. During early G1, we found three Cohesin-STAG2 molecules to be on average bound for 120 seconds per megabase of DNA. Assuming they extrude loops with the estimated rate of 1 kb/s (Kim et al., 2019; Davidson et al., 2019), they would form 120 kb large loops and would thus be relatively unlikely to encounter each other within one megabase (Table 1). In late G1 however, about 8 Cohesin-STAG2 complexes are bound per megabase with a similar residence time (Table 2), making Cohesin-STAG2 encounters between eight 120 kb sized loops within one megabase much more likely. The fact that we observe a quantitative shift in the intensities of the Cohesin-STAG2 spot distribution from early to late G1 (Fig. 5D) in fact suggests that Cohesin complexes not only encounter each other transiently, but potentially stay associated with each other when they meet, which would induce stacking and nesting of loops. To test if such nested and stacked loops indeed form in late G1 in single cells in a Cohesin-STAG2 dependent manner, we again made use of our nanoscale DNA tracing of interphase cells targeting the same three 1.2 megabase TAD regions as before. The 3D folds of these regions indeed revealed stronger nesting and stronger compaction of these regions compared to early G1, and again showed that this is largely dependent on Cohesin-STAG2 (Fig. 5E-H, Suppl. Fig. SJ-0). In conclusion, due to its continuous nuclear import, Cohesin-STAG2 crosses a critical occupancy threshold on the genome within the first 1-2 hours after mitosis that leads to a high probability of encounters between Cohesin-STAG2 complexes, accompanied by increased formation of nested loops inside TAD-scale compact domains of the interphase genome.

**Table 2.** Quantitative data summary: G1.

| POI | G1 (= 2-4h past AO) |  |  |  |  |  |  |  |  |
| --- | --- | --- | --- | --- | --- | --- | --- | --- | --- |
|  | nuclear, relative to 2h post AO | absolute number in nucleus | % bound (spot-bleach, average), interphase time-point | % bound (FRAP), G1 timepoint | Residence time, dynamically bound pool | absolute number, bound | absolute, chromatin bound, per Mb | fraction long term bound, per Mb | absolute, long term bound, per Mb |
| SMC4 | 100% | 174000 | 0.1 |  |  | 17400 |  |  |  |
| NCAPH | 100% | 125000 | 0.09 |  |  | 11250 |  |  |  |
| NCAPH2 | 100% | 25600 | 0.02 |  |  | 512 |  |  |  |
| CTCF | 100% | 124000 | 0.72 | 0.72 | 139 | 89280 | 5.7 | 0.488 | 2.8 |
| RAD21 | 100% | 238000 | 0.59 | 0.77 | 172 | 183260 | 11.6 | 0.277 | 3.2 |
| STAG1 | 100% | 53500 | 0.59 | 0.76 | 276 | 40660 | 2.6 | 0.39 | 1.0 |
| STAG2 | 100% | 183000 | 0.5 | 0.71 | 126 | 129930 | 8.2 | 0.107 | 0.9 |

### A double hierarchical loop model quantitatively explains the transition from mitotic to interphase loop extruder driven genome organization

0ur systematic, quantitative time-resolved mapping of mitotic and interphase loop extruders in single cells shows that the interphase genome is sequentially organized into compact TAD-scale regions which then compact further by internal stacking of nested loops. This is highly reminiscent of the previously proposed nested loop organization in mitosis (Walther et al., 2018), in which chromosomes are organized by the sequential action of two Condensin complexes (Walther et al., 2018; Gibcus et al., 2018). In this model of establishing mitotic architecture, the less abundant Condensin II loop extrusion motor first forms large DNA loops during prophase that become subsequently nested by the more abundant Condensin I, once it gains access to chromosomes during prometaphase.

0ur study now shows, that following chromosome segregation and nuclear envelope reformation, some Condensins are still bound to chromatin, while Cohesins and CTCF are rapidly imported into the newly formed nucleus, leading to a co-occupancy of the genome by 3 Condensins and 3 Cohesins per megabase of DNA in early G1. Very interestingly, when the interphase loop extruders Cohesins start binding the genome, they do so independently of Condensins, but like Condensins during mitotic entry also in a sequential manner during mitotic exit. First, the rapid and synchronous nuclear import of Cohesin-STAG1 and CTCF (completed within l0 minutes after A0) and their immediate chromatin-binding at relatively low abundance (3 complexes bound per megabase) with a long residence time (4 minutes) builds up the first compact interphase structures even in the absence of Cohesin-STAG2. In a second step, the slowly imported Cohesin-STAG2 (complete only 2 hours after mitosis) then binds in higher abundance (8 complexes bound per megabase) and with a shorter residence time (2 minutes), leading to the generation of many smaller loops (∼120 kb), and frequent encounters and likely stalling with neighboring Cohesin-STAG2 complexes, leading to stacking of nested loops inside the larger STAG1 defined domains. We therefore propose a double hierarchical loop model for the transition from mitotic to interphase loop extruder driven genome architecture, in which the Condensin-based, randomly positioned nested loop architecture established during mitotic entry is replaced by a less compact, but conceptually similar Cohesin-driven nested loop architecture, positioned by CTCF, from mitotic exit to early G1 (Fig. 6).

**Figure 6.**
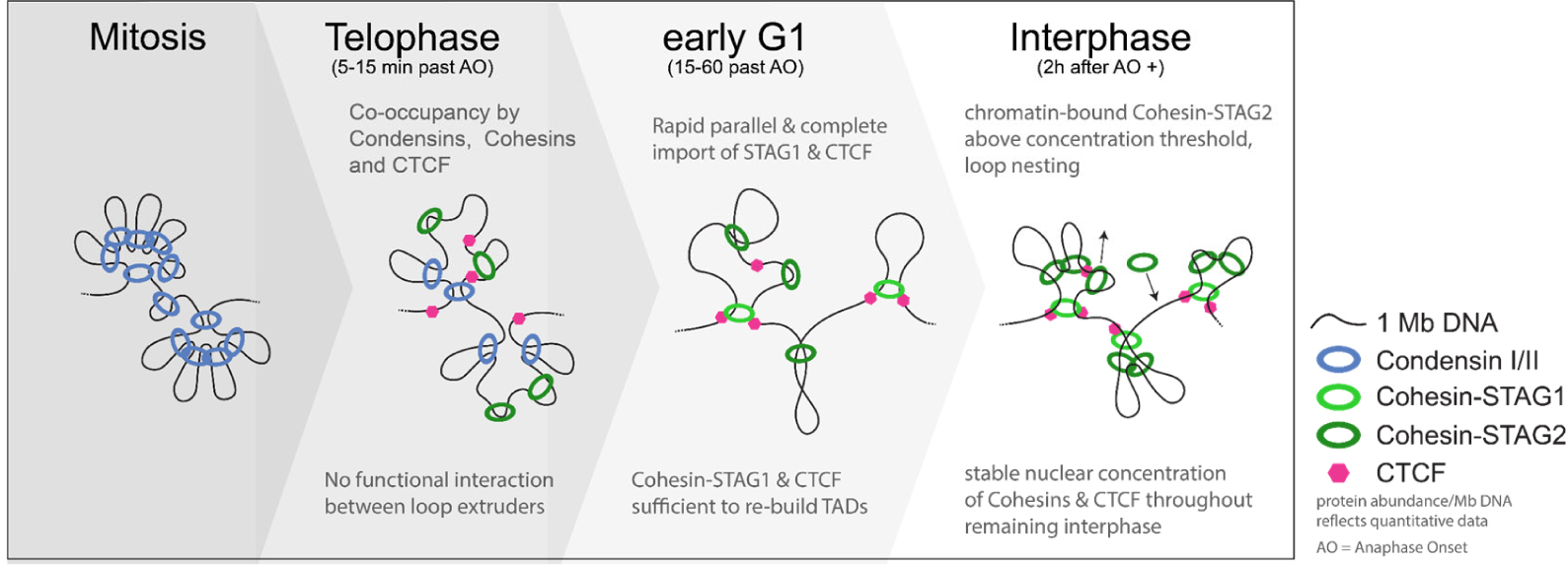
A new model for the mitosis-to-interphase transition of genome organization by loop extrusion. Mitotic chromosomes are majorly organized by the abundant Condensin complexes (l2/Mb, of those ∼l.S are Condensin II), only a small fraction of CTCF (about l4,000 proteins, ∼ l0% of cellular pool) binds chromatin very transiently and a small fraction of the cellular Cohesin holds together sister chromatids until anaphase. During telophase and early Gl, not all Condensins dissociate right away, leading to a co-occupancy of 3 Condensins and 3 Cohesins per megabase DNA and a significant pool of Condensin I that remains nuclear until the next mitosis starts. Cohesin-STAGl and CTCF are fully imported into the newly formed nucleus within l0 minutes after anaphase onset and are sufficient to build interphase TAD structures in the absence of Cohesin-STAG2. Cohesin-STAG2 only completes its import 2 hours after anaphase onset and - due to its abundance on chromatin upon full import (8 complexes per megabase) - frequently crashes into neighboring loop extruders, leading to increased nesting of sub-TAD loops. While Cohesin-STAGl is stabilized through CTCF and SMC3 acetylation (Wutz et al., 2020), CTCF is increasingly stabilized on chromatin through its association with Cohesin on chromatin. After completed import of Cohesins and CTCF, their nuclear concentrations remain stable irrespective of the DNA content of the nucleus.

## Discussion

This study provides comprehensive quantitative and time-resolved data on the chromatin-binding of Condensins and Cohesins throughout mitotic exit and G1. In addition to the new data it provides, it furthermore allows to integrate many previous more qualitative and individual observations into an overall, internally consistent and quantitative model of how the loop extruder-based genome organization is handed over from mitosis to interphase.

### A role for chromatin-bound Condensin during telophase?

The Condensin-driven mitotic chromosome organization, previously proposed to be best explained by an axial arrangement of nested DNA loops (Gibcus et al., 2018; Walther et al., 2018), is rapidly lost during telophase when 75% of Condensins unbind DNA. Consistent with a previous report (Abramo et al., 2019), we find that this rapid removal of Condensins is followed by import of Cohesin and CTCF into the newly forming nuclei, leading to a co-occupancy of only 3 Condensins and Cohesins per megabase during early G1, which we show is the lowest number of genome-associated loop extrusion complexes at any time during the cell cycle. Nonetheless a significant fraction of Condensins remains chromatin bound during telophase and early G1, leading to a so far unappreciated pool of Condensin I to be retained in the nucleus during interphase. However, in interphase nuclear Condensin I is unlikely to be actively engaged in processive loop extrusion due to its mitosis-specific loading onto chromosomes (Hirano et al., 1997) regulated via phosphorylation of the NCAPH N-terminal tail (Tane et al., 2022) and mitotic activation by KIF4A (Cutts et al., 2024). However, it could be that the retained fraction of Condensins has a transient role during telophase and early G1 to facilitate the removal of intra-chromosomal catenations as suggested recently (Hildebrand et al., 2024).

### The two Cohesin isoforms bind sequentially and likely have different structural roles

Consistent with previous systems analysis of mitotic protein networks (Cai et al., 2018), we found that Cohesin-STAG1 and CTCF are rapidly imported into the newly formed daughter cell nuclei after cell division and are sufficient in numbers and loop extrusion processivity to form the first interphase TAD-scale loops shortly after mitosis. By contrast, we found that the more abundant second Cohesin isoform STAG2 is imported slowly over the course of 2 hours and is dispensable for the generation of these TAD-scale compact structures in early G1 as well as later in interphase. Due to its later binding, its higher abundance and its short residence time on chromatin, Cohesin-STAG2 leads to shorter and nested loops within the already established larger Cohesin-STAG1 loops. We speculate that this STAG2-dependent highly dynamic sub-structuring of the more stable TAD loops could promote cell-type-specific intra-TAD contacts between enhancers and promoters independently of CTCF, explaining cell-type dependent effects of Cohesin-STAG2 mutations or depletion (Kojic et al., 2018; Viny et al., 2019). While we found that the overall fraction of chromatin-bound proteins increases for both Cohesin isoforms and CTCF similarly from early to later G1, Cohesin-STAGl and CTCF became specifically stabilized on chromatin during Gl. 0ur observations are consistent with recent reports that Cohesin-STAG1 is stabilized through CTCF binding and acetylation of its SMC3 subunit by Esco1 (Wutz et al., 2020), suggesting a continued role of Cohesin-STAG1 in the generation and maintenance of long-range loops during interphase that are further sub-structured by Cohesin-STAG2.

### Chromatin binding of CTCF is stabilized by Cohesin

0ur FRAP analysis early after mitosis and later during G1 enabled us to see a clear increase in the fraction of stably chromatin bound CTCF. Interestingly, this stabilization was dependent on the presence on Cohesin, and occurred progressively throughout G1 when we found both Cohesin isoforms to increasingly co-localize with CTCF. It is therefore likely that CTCF’s stabilization is due to interaction with one or both chromatin-bound Cohesin isoforms, which may serve as additional anchors at CTCF-sites. Combined with the fact that Cohesin-STAG1 is preferentially associating with CTCF (Kojic et al., 2018) and is stabilized in part through CTCF (Wutz et al., 2020) our data suggests that Cohesin-STAG1 and CTCF mutually stabilize each other on chromatin, which may be important to stabilize longer lived loop structures in interphase, equivalent of TADs.

### The oligomerization state of Cohesin

Using structured illumination microscopy of the isoform-shared Cohesin subunit RAD21, it was recently reported that the majority of the loop-extruding Cohesin is present in dimers or multimers (0chs et al., 2024). Consistent with this finding, our STED super-resolution imaging of isoform-specific Cohesin subunits revealed that the less abundant Cohesin-STAG1 is present as a monomer, but that the more abundant isoform Cohesin-STAG2 undergoes dimerization on chromatin in later G1 in a concentration dependent manner. In addition, we found Cohesin-STAG2 to be on average bound to chromatin for 120 seconds and increasing its occupancy from 3 to 8 complexes per megabase from early to late Gl. 0ur data thus quantitatively explains how Cohesin STAG2 dimers form, if we assume the bound complexes extrude loops with the reported speeds (i.e. 0.5-2 kb/s, Kim et al., 2019; Davidson et al., 2019): While 3 randomly loaded Cohesin-STAG2 complexes per megabase DNA are very unlikely to encounter each other during early G1 due to the relatively small loops they can make during 2 min (around 120 kb), encounters and potential stalling between loops of this size become much more likely when Cohesin-STAG2 is fully imported and present at 8 copies per megabase in late G1. While we cannot exclude that Cohesin-STAG2 dimers continue active loop extrusion, our data would be consistent with the view that the default state of Cohesin complexes in loop extrusion is monomeric and that dimers result from encounters and potential stalling events.

### A comprehensive and quantitative dataset to constrain next generation polymer models

In summary, our systematic and quantitative assessment of Condensin and Cohesin loop extruder dynamics on chromatin provides a comprehensive and integrated view of the transition from mitotic to interphase genome organization. Given the sequential import of Cohesin isoforms, their chromatin binding dynamics, their different abundance as well as impact on chromatin upon depletion, we propose a hierarchical nested loop model for the establishment of the interphase genome organization by the Cohesin loop extruders after mitosis. 0ur model is conceptually similar to the hierarchical nested loop architecture proposed for the establishment of Condensin driven mitotic organization (Gibcus et al., 2018; Walther et al., 2018). While the sub-structuring of large Condensin II loops in mitosis by Condensin I serves to laterally compact mitotic chromatin and confers additional mechanical rigidity to chromosomes (0no et al., 2003; Shintomi and Hirano, 20ll; Green et al., 2012; Houlard et al., 20lS), we think that the sub-structuring of large STAG1 loops (Kojic et al., 2018; Wutz et al., 2020) by STAG2 aids TAD-scale compaction and specific intra-TAD contact enrichment, potentially in a cell type and species-specific manner (Dixon et al., 2012; Phillips-Cremins et al., 2013; Rao et al., 2014). The quantitative data and understanding provided by our study should provide a comprehensive quantitative basis for next generation predictive and mechanistically explanatory models of genome organization.

## Supporting information

Antibodies

Cell Lines

gRNAs

## Acknowledgements

We thank the EMBL Advanced Light Microscopy Facility (ALMF) and the EMBL Imaging Center for microscope support as well as the Centre for Bioimage Analysis for support related to image analysis. We thank Gordana Wutz for valuable discussions and supply with antibodies and cell lines related to Cohesin and CTCF. This work was supported by grants from the National Institutes of Health Common Fund 4D Nucleome Program (Grant U0l EB02l223 / U0l DA047728) to J.E., as well as by the The Paul G. Allen Frontiers Group through the Allen Distinguished Investigator Program to J.E.. A.B. has received a PhD fellowship from the Boehringer Ingelheim Fonds and K.S.B. was supported by the Alexander von Humboldt foundation. Work in the laboratory of J.-M.P. has received funding from Boehringer Ingelheim, the Austrian Research Promotion Agency (Headquarter grant FFG-852936), the European Research Council (ERC) under the European Union’s Horizon 2020 research and innovation programme (grant agreements No 693949 and No l0l020SS8), the Human Frontier Science Program (grant RGP00S7/20l8) and the Vienna Science and Technology Fund (grant LS19-029).

## Author contributions

A.B. and J.E. conceived the project. K.S.B and J.E. supervised the project. A.B. designed, performed and analyzed all experiments with technical support from N.R.M. related to spot-bleach data acquisition, from K.S.B. related to FCS-calibrated imaging, FRAP and chromatin trace analysis, from M.J.H. related to the segmentation of live-cell imaging data, from M.L. related to STED image acquisition, from H.P. related to protein degradation, and from A.H. related to automated live-cell imaging. N.R.M. and W.Z. generated and validated genome-edited cell lines created within this study. A.B. and J.E. wrote the manuscript with reviewing and editing performed by all authors. Funding was acquired by A.B., K.S.B., J.E. and J.-M.P.

## Materials and Methods

### Cell Culture

HeLa Kyoto cells (RRID: CVCL_l922) were obtained from S. Narumiya (Kyoto University, Kyoto, Japan) and cultured in high-glucose DMEM (4l96S-062, Thermo Fisher Scientific) supplemented with 10% FBS (l0270-l06, Lot. 42F2388K, Thermo Fisher Scientific), 100 U/ml penicillin-streptomycin (lSl40-l22, Thermo Fisher Scientific) and 1 mM sodium pyruvate (ll360-039, Thermo Fisher Scientific) at 37°C, S% C0_2_ unless otherwise stated. Cells were grown in cell culture dishes (Falcon) and passaged every 2-3 days via trypsinization with 0.05% Trypsin-EDTA (2S300-0S4, Thermo Fisher Scientific) at 80-90% confluency. Mycoplasma contamination was checked regularly and confirmed negative.

### FCS-calibrated confocal time-lapse imaging

Cell samples for FCS-calibrated confocal time-lapse imaging were prepared according to Politi et al. (2018). Specifically, two days before the experiment, two 0.34 cm^2^ wells of an 18-well chambered coverglass (Ibidi µ-slide, 8l8l7) were seeded with 3750 HK WT cells. 24 hours prior to the experiment, one well of HK WT cells was transfected with a plasmid expressing monomeric EGFP and 2,000-4,000 genome-edited cells expressing the protein of interest (P0I) endogenously tagged with (m)EGFP were seeded in a third well. l.S hours prior to imaging, DMEM medium was exchanged to phenol-red free C0_2_-independent imaging medium based on Minimum Essential Medium (Sigma-Aldrich, M3024) containing 30 mM HEPES (pH 7.4), 10% FBS, 1X MEM non-essential amino-acids (lll40-0S0, Thermo Fisher Scientific) and S0-l00 nM S-SiR-Hoechst (gift from G. Lukinavičius, Bucevičius et al., 2019). In addition, after l.S hours of DNA-labelling by 5-SiR-Hoechst, 500-kDa dextran-Dy481XL (Cai et al., 2018) was added to the genome-edited cells to facilitate cell segmentation.

Fluorescence Correlation Spectroscopy (FCS)-calibrated imaging was performed on Zeiss LSM780 (equipped with ConfoCor 3 unit, controlled by ZEN 2.3 Black software, Version l4.0.l8.20l, Zeiss) and LSM880 (controlled by ZEN 2.1 Black software, Version l4.0.9.20l, Zeiss) laser-scanning microscopes with an inverted Axio 0bserver microscope stand, equipped with an in-house constructed incubation chamber for temperature control set to 37°C (without C0_2_ due to use of C0_2_-independent imaging medium) and using a C-Apochromat 40x/1.2 W Korr UV-Vis-IR water-immersion objective (42l767-997l-7ll, Zeiss). Microscope calibration by FCS was performed as described by Politi et al. (2018), but using 10 nM Atto488 carboxylic acid (AD 488-2l, ATT0-TEC, Kapusta, 2010) in ddH_2_0 instead of AF488 coupled to a H20-hydrolyzable NHS ester group to estimate the confocal volume in FCS measurements. This led to ∼30% larger confocal volume estimates in better agreement with other methods for confocal volume determination (Buschmann et al., 2009). This change resulted in a systematic drop of the protein concentrations measured proportional to the change in confocal volume size, compared to previous measurements using AF488-NHS (Politi et al., 2018). Ten FCS-measurements of l minute each were performed to estimate the effective confocal volume in the well with Atto488 solution. FCS-measurements of 30 seconds were performed in the nucleus and cytoplasm in WT cells not expressing mEGFP to determine background fluorescence and photon counts. Experiment-specific calibration factors were obtained from interphase cells expressing mEGFP by correlating measured fluorescence intensities and absolute mEGFP concentration calculated from 30 seconds FCS-measurements (Politi et al., 2018).

Calibrated 4-dimensional confocal time-lapse imaging was performed on cells expressing the mEGFP-tagged protein of interest (P0I) using a combination of MyPic macros for ZenBlack software (https://git.embl.de/grp-ellenberg/mypic), AutoMicTools library (https://git.embl.de/halavaty/AutoMicTools) for ImageJ (Schindelin et al., 2012) and ilastik (Berg et al., 2019). Specifically, metaphase cells were automatically identified in multiple pre-defined fields of view by low-resolution imaging of the DNA channel (5-SiR-Hoechst). Subsequently, cells of interest were imaged for the next lS0 minutes with a time-resolution of 2 min to capture anaphase onset (A0) and l20 minutes of progression through mitotic exit with 31 z-slices with a voxel size of 250 nm in xy and 750 nm in z, covering a total of 7Sx7S µm in xy (300×300 pixels) and 22.5 µm in z, which was sufficient to cover the whole cell volume, in the GFP ((m)EGFP-tagged P0I), DNA, Dextran-Dy481XL (extracellular space), and transmission channels. A previously developed computational pipeline (Cai et al., 2018) was adapted to track and segment dividing cells from high-zoom time lapses in 3D based on the nuclear (SiR-Hoechst) and cellular (Dextran-Dy481XL) landmarks. The third eigenvalue of the segmented chromatin mass, representing the thickness of the chromosomal volume, was utilized to detect A0 as chromosomes begin to be segregated towards opposite cell poles. All mitotic exit time-series were aligned to A0 and set as the t=0 min timepoint. All individual aligned time-series displayed a very consistent increase in chromatin volume over time, rendering any further alignment dispensable.

### Estimation of protein numbers from FCS-calibrated images

Fluorescence intensities in image voxels were converted to absolute protein concentrations and numbers based on the experiment-specific calibration line (calibration factor (= slope) and background intensity) and the 3D binary masks of nucleus and the cell. The average protein concentration was calculated by multiplying the calibration factor (slope of the calibration line) to the average background corrected fluorescent intensity in all nuclear, cellular or cytosolic pixels (cytosol = within the cell, but excluding the nucleus). The absolute protein number inside each compartment was achieved by integrating all background-corrected fluorescent intensities and multiplying them with the calibration factor.

### Full Cell Cycle Imaging

About 7S0-l000 genome-edited cells expressing the P0I endogenously tagged with EGFP were seeded two days before the experiment into a 0.34 cm^2^ well of an 18-well chambered cover glass (Ibidi µ-slide, 8l8l7) and incubated at 37°C, S% C0_2_. 20 hours day later, cells were arrested in S-phase for lS-l6 hours by changing the medium to DMEM supplemented with 2 mM thymidine (Tl89S, Sigma). Cells were subsequently released from S-phase arrest by washing 3 times with DMEM. 4 hours after release, medium was exchanged to phenol-red free, C0_2_-independent imaging medium (see above) containing S0-l00 nM S-SiR-Hoechst and one hour later 500-kDa dextran-Dy481XL was added as a cell outline marker (added later due to interference with efficient SiR-Hoechst staining). Imaging was started 6 hours after release from S-phase, well before the first mitotic division. As a control of the effect of S-phase arrest, ∼3750 asynchronous cells were seeded one day before imaging into a well of an 18-well Ibidi µ-slide and imaging was carried out l.S hours after addition of imaging medium containing S-SiR-Hoechst and addition of 500-kDa dextran-Dy481XL. Imaging was carried out on a Zeiss LSM780 and LSM880 using a C-Apochromat 40x/1.2 W Korr UV-Vis-IR water-immersion objective (42l767-997l-7ll, Zeiss) with a custom-made objective cap for automated water dispension, with a field of view (F0V) size of l77.l2xl77.l2 µm covering a z-range of 22.5 µm with 253 nm pixel size in xy and 750 nm in z and a pixel dwell time of 0.76 µsec. 0.2% laser power of the 488 nm Argon laser line was used to ensure minimal bleaching and GFP fluorescence was recorded on the GaAsP detector (499 nm-SS3 nm range, gain set to ll00). 4 F0V were automatically imaged every 10 minutes with an autofocus step before every single 3D stack (based on peak reflection of Sl4 nm laser line at glass-sample interface). Depending on the cell cycle length and whether synchronous or asynchronous cells were used, total imaging time varied from 25 to 40 hours, in order to capture two subsequent mitosis events for most cells present in the F0V. Image data was processed using an adapted computational pipeline (Cai et al., 2018) performing 3D segmentation based on chromatin (5-SiR-Hoechst) and cellular landmarks (500 kDa Dextran), as well as cell tracking of single cells using the 3D centroid of the chromatin mass. After manually filtering out duplicate or poorly segmented single cell tracks, single cell cycles were cropped out based on the cellular and nuclear volume information, resulting in a list of full cell cycle tracks ranging from one anaphase/telophase to the next. These full cell cycle tracks were aligned to the first division and subsequently interpolated and fit to a common average cell cycle timing. Calibration of the measured fluorescent intensities was performed not through direct FCS-calibrated imaging, but by setting the number of proteins inside a cell (N_cell) in the second mitosis (when the S-phase arrest effect has ceased) to the mean number of proteins inside a cell measured in asynchronous FCS-calibrated metaphase cells, resulting in a conversion factor that was used to transform measured fluorescent intensities to absolute protein numbers and concentrations at all other timepoints. While bleaching of GFP-tagged proteins was not tested over the course of an entire cell cycle, we assume it to be minimal due low laser exposure (488 nm: 0.2%, pixel dwell: 0.76 µsec, l stack every l0 min) and the fact that cellular concentrations of all proteins did not change from one mitosis to the next.

### Simple Western

Protein separation, immunodetection and quantification from cell lysates was performed in a Jess Automated Western Blot System (Bio-Techne), using 12-230 kDa and 66-440 kDa Fluorescence separation capillary cartridges (SM-FL004-1, SM-FL00S-l, Bio-Techne). For this, total protein lysates were prepared for each cell line and condition of interest by growing cells in a 10-cm until ∼80% confluency, subsequently washing with PBS and resuspending cells in 500 μl of lysis buffer (RIPA buffer (R0278, Sigma-Aldrich), 1 mM PMSF (P7626, Sigma-Aldrich), c0mplete™ EDTA-free Protease Inhibitor Cocktail (04693l3200l, Roche, l tablet/l0 ml) and PhosST0P (490684S00l, Roche, l tablet/l0 ml)) with the help of a cell scraper (on ice). Cells were then lysed by two cycles of freezing in liquid nitrogen and thawing at 37 °C. After centrifugation for 10 min at ∼l6,000xg, 4°C, the supernatant containing soluble total protein extracts was separated and kept at −80°C until use. Total protein was quantified with a Pierce BCA Protein Assay Kit (23227, Thermo Fisher Scientific) and diluted to 0.4 µg/µL final concentration including 1x Master Mix (from EZ Standard Pack 1 (PS-ST01EZ-8, Bio-Techne). Loading of samples and detection reagents into the Simple Western (SW) microplate was conducted following the provider’s instructions. Detection was achieved by ECL using anti-rabbit and anti-mouse secondary HRP antibodies (042-206/ 042-205, Bio-Techne) and Luminol-S/Peroxide solution (043-3ll/043-379, Bio-Techne). Capillary electrophoresis run and analysis was conducted with the Compass for SW software (Bio-Techne) following the provider’s guidelines.

### Preparation of homozygous endogenous knock-in cell lines

Genome-edited cell lines generated in this study (HK Rad21-EGFP-AID CTCF-Halo-3xALFA #C7 and HK NuplS3-mEGFP-FKBPl2^F36V^ #C10 (dTAG technology: Nabet et al., 2018) were obtained by C-terminal tagging of CTCF and NuplS3 in HK RAD21-EGFP-AID (Davidson et al., 2016) or HK WT parental cell lines, respectively, using the CRISPR/Cas9 method. In brief, a linear DNA donor sequence encoding for the tag of interest (and corresponding 50 base pair long homology arms) was electroporated into the parental cell line, together with the catalytic Cas9/gRNA ribonucleoparticle complex, as previously described (Koch et al., 2018; Kueblbeck et al., 2021 *Preprint*). For this, we used Alt-R™ S.p. HiFi Cas9 Nuclease V3 (l08l06l, IDT) and single gRNAs (see Supplementary Information). Edited cells expressing the tags of interest were selected by FACS sorting and the correct tagging of all target copies was subsequently validated as described in (Kueblbeck et al., 2021). Expression of the tagged protein of interest (P0I) at endogenous levels was confirmed by simple western and confocal microscopy, the latter also indicating correct subcellular localization of the P0I. Homozygous tagging of the P0I was confirmed by PCR screening, simple western and digital PCR. Digital PCR (dPCR) allows to quantify the copy number of specific sequences of interest in a template genome, by partitioning the amplification reaction (including a primer pair and an internal fluorescent probe, per region to be quantified) into thousands of nanodroplets, each containing 0-few DNA molecules. Upon amplification of the region of interest in a given droplet, the specific internal probe is released from the DNA and fluorescence is detected. The count of fluorescent vs non-fluorescent droplets is read out and used to quantify the absolute amount of template DNA. The triple-color dPCR assay used in this work allowed us to quantify: the total number of tags (“allGFP” or “allHalo”) integrated into the genome, the number of tags inserted at the intended target locus (“HDR”, homologous-directed repair after Cas9-directed DNA cut) and the copy number of a reference sequence located in the vicinity of the target locus. This setup therefore allows to quantify how many endogenous alleles are tagged, as well as the detection of excess off-target tag integrations within the recipient genome. Finally, the correct sequence and positioning of the integrated tags was corroborated by PCR-amplification and sequencing of the edited genomic regions.

### Fluorescence recovery after photobleaching

Cells for FRAP measurements were seeded at a density of 2.Sxl0^S^ cells/ml into Ibidi glass bottom µ-Slide channels (80607, Ibidi) one day prior to imaging. DMEM was replaced by C0_2_-independent imaging medium (as above) containing S0-l00 nM S-SiR-Hoechst at least 1 hour before imaging. FRAP experiments were performed on a LSM880 laser-scanning microscope with an inverted Axio 0bserver controlled by ZEN 2.1 Black software (Version l4.0.9.20l, Zeiss), equipped with an in-house constructed incubation chamber for temperature control set to 37°C and using a C-Apochromat 40x/1.2 W Korr UV-Vis-IR water-immersion objective (42l767-997l-7ll, Zeiss). Cells in metaphase and early G1 were selected manually based on their chromatin staining and FRAP of metaphase cells was performed as described previously (Walther et al., 2018). Cells in G1 stage were selected manually based on nuclear size and filtered out computationally based on a nuclear size threshold of less than l0S0 µm^3^ corresponding to the size of cells about 5 hours into the cell cycle according to full cell cycle data of asynchronous cells (exact nuclear size was derived from a 3D stack covering the whole chromatin mass, segmented with a previously developed script (Cattoglio et al., 2019). A single image was recorded prior to bleaching, recording 5 z-planes in metaphase and early G1, 3 z-planes in G1 with a pixel size of 2l3×2l3×7S0 nm, pixel dwell 1.7 µsec and a F0V size of 27.2Sx27.2S µm for metaphase and G1 cells and of 42.Sx42.S µm for early G1 cells, respectively in the EGFP (488 nm argon laser line, excitation power: 1%, GaAsP detection range set to 499 nm - 562 nm, gain set to 1000) and SiR-Hoechst channels (633 nm diode laser, excitation power 0.2-0.4%, GaAsP detection range set to 641 nm - 696 nm, gain set to 1000). Subsequently, a square region covering half of the chromatin / nucleus area in the middle z-plane was bleached using similar laser power for metaphase, early Gl and Gl cells (488 nm laser power: 100%). While metaphase plates were bleached with one bleach step (4S x 3S pixels, lS0 repetitions), early Gl and Gl cells were bleached 3 times within 30 seconds to completely bleach the freely diffusion soluble pool (4S x 3S pixels for eGl, 60 x S0 pixels for Gl, 3x 50 repetitions), enabling the determination of chromatin-bound fractions. The fluorescent recovery was recorded by time-lapse imaging every 20 seconds for another 30 frames with the settings described for the pre-bleach image, resulting in minimal bleaching throughout the imaging period (<10%).

FRAP image analysis was performed using a previously developed custom-written ImageJ script (Walther et al., 2018), adapted to enable the analysis of metaphase, early Gl and Gl cells at the same time, as well as an R-script for downstream data processing (Walther et al., 2018). In brief, this analysis script aggregates the (m)EGFP-P0I and SiR-Hoechst fluorescence intensity data along the major 2D chromatin axis (segmented using SiR-Hoechst channel) into a 1D profile. Using a gap of 14 pixels in the center of the 1D profile, the border of the bleaching R0I was omitted to avoid boundary effects. The weighted mean fluorescence intensities (using SiR-Hoechst) in the unbleached and bleached regions were computed as described in (Walther et al., 2018). As in (Gerlich et al., 2006; Walther et al., 2018), the weighted normalized difference between the unbleached and bleached region

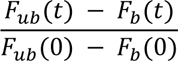

was used as a readout for the residence time and immobile fraction. A single exponential function

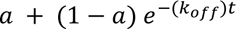

was employed to fit the normalized fluorescence recovery data. The parameter *a* represents the immobile fraction and k_off_ is the unbinding rate constant.

### FRAP to investigate Cohesin-dependence of CTCF chromatin association

FRAP measurements of CTCF after depletion of RAD21 were carried out in G1 cells of genome-edited HK cells in which all alleles of RAD21 were tagged with an AID degron and EGFP and all alleles of CTCF were tagged with Halo (see above). G1 cells were selected based on nuclear volume, but no stringent size filter was applied since the variance of individual measurements was found to be minimal and not dependent on nuclear volume. Complete depletion of RAD21 in these genome edited cells was achieved by incubation with Inole-3-acetic acid (IAA, ISl48, Sigma) for at least l.S hours. For rescue of RAD21 depletion, exogenous RAD21-EGFP was overexpressed for at least 24 hours prior to the start of the experiment. FRAP measurements were carried out as described above, however bleaching and imaging of fluorescence recovery was performed using 561 nm excitation of the Halo-TMR (G8252, Promega) ligand coupled to endogenous CTCF-Halo (excitation power: 0.7%, GaAsP detection range set to 570-624, gain set to l000) after l0 minutes of labelling with Halo-TMR at a concentration of 100 nM at 37°C in imaging medium. Interestingly, we found that CTCF-Halo displayed a reduced chromatin residence time and immobile fraction in the absence of IAA, unlike CTCF-EGFP endogenously tagged in a different cell line. We found that this correlated with a leaky degradation of RAD21 in the RAD21-EGFP-AID CTCF-Halo cell line, reducing RAD21 levels about 40% relative to our CTCF-EGFP line (using Simple Western of asynchronous cell lysates, RAD21 detected via anti-RAD2l antibody (0S-908, Merck Millipore, l:S0, Suppl. Fig. 4G). 0verexpression of RAD2l rescued this effect, bringing CTCF-Halo residence time and bound fraction almost back to WT levels (data not shown). For comparison with our ΔRAD21 and ΔRAD21+rescue conditions, we therefore decided to use our CTCF-measurements as WT reference condition.

### Cell synchronization by mitotic shake-off

To synchronize HK cells in mitosis for subsequent protein degradation or timed release into early Gl or Gl, we used a combination of Nocodazole treatment and a mitotic shake-off. In brief, cells were regularly passaged (every second day) and seeded into a T-l7S flask (3S3ll2, Corning) to reach a confluency of around 80% after 16-24 hours of incubation. 0ne hour prior to mitotic shake-off, cells were incubated in 12 mL of DMEM complete medium supplemented with 82 nM Nocodazole (SMLl66S, Sigma-Aldrich) to enrich mitotic cells. The mitotic shake-off was conducted by banging 5 times the cell culture flask on a table covered with ∼5 paper tissues. After confirming the detachment of most mitotic cells by inspection on a microscope, the mitotic cell suspension was transferred to a lS mL Falcon tube and centrifuged for 3 minutes at 90xg. The resulting cell pellet was resuspended in lS0 µL DMEM + 82 nM Nocodazole and the cell density was counted. 35 µL of cells at a desired density (between l.2xl0^6^ cells/ml and 2.Sxl0^6^ cells/ml) were seeded into an Ibidi µ-Slide glass bottom slide (80607, Ibidi) with channels pre-coated for lS minutes with poly-L-lysine (P8920, Sigma). Ibidi slides were incubated for lS minutes at 37°C, S% C0_2_ to allow cells to attach. 100 µL of DMEM complete medium supplemented with 82 nM Nocodazole was added to cells in every Ibidi µ-Slide channel prior to any further treatment.

### Immunofluorescence

Fixed cells were prepared for immunostaining by permeabilization with 0.25% Tergitol (lSS9, Sigma) in PBS for lS minutes and subsequent incubation in blocking buffer (2% BSA, 0.05% Tergitol in PBS) for at least 30 minutes at room temperature (RT, 20-25°C in this work). Primary antibody incubation was performed in blocking buffer at 4°C in a humidified chamber overnight (16-24 hours), followed by washing with blocking buffer (3 times, 5 min). Secondary antibody hybridization was performed in blocking buffer for 1h at RT. After washing with PBS (3 times, 5 min), samples were post-fixed with 2.4% PFA (lS7l0, EMS) in PBS for lS minutes, quenched with 100 mM NH_4_Cl in PBS for 10 minutes and washed in PBS. Samples used for LoopTrace-based chromatin tracing were permeabilized with Triton X-100 instead of Tergitol at the same concentration for consistency with previous experiments.

### Protein depletion during mitosis

For the degradation of NuplS3, SMC4, RAD21 and CTCF during mitosis, we used genome-edited HK cells in which all copies of the P0I were endogenously tagged with a dTAG degron system (NuplS3-mEGFP-FKBPl2^F36V^, Nabet et al., 2018, 2020), or an Auxin-inducible degron tag (SMC4-Halo-mAID (Schneider et al., 2022), RAD21-EGFP-AID (Davidson et al., 2016), CTCF-mEGFP-AID (Wutz et al., 2017)). Nocodazole-assisted mitotic shake-off was conducted as described above and 35 µL of mitotic cells were seeded into Ibidi glass bottom µ-slides (80607, Ibidi) pre-coated with poly-L-lysine (lS minutes) at a density of 2-2.Sxl0^6^ cells/ml. Cells were allowed to attach for lS minutes at 37°C, S% C0_2_. Subsequently, the depletion of degron-tagged proteins was conducted for l.S hours in the presence of 82.5 nM Nocodazole and each specific degradation-triggering ligand (NuplS3: 2S0 nM dTAG-l3 (SML260l, Sigma) & S00 nM dTAG^V^-l (69l4, Tocris); SMC4: l uM S-Ph-IAA (30-003, BioAcademia); RAD21 & CTCF: S00 uM IAA). Afterwards, cells were released into mitotic exit by washing out Nocodazole through cell incubation for 45-90 minutes in fresh medium supplemented with dTAGs, 5-Ph-IAA or IAA, respectively. Then, cells were either pre-extracted by washing in PBS and then incubating with 0.25% Tergitol in PBS for 1 minute followed by PFA-fixation (ΔSMC4, ΔNIPBL), or fixed directly with 2.4% PFA in PBS for lS minutes (ΔNuplS3), followed by quenching of PFA with 100 mM NH_4_Cl in PBS and washing with PBS. Immunofluorescence was performed as described above, using the following primary antibodies: mouse anti-RAD2l (0S-908, Merck Millipore, l:S00), rabbit anti-SMC2 (abl04l2, Abcam, l:l000); rabbit anti-CTCF (07-729, Merck Millipore, 1:2000) or rabbit-anti CTCF (Wutz et al., 2020, Glycine Elution, l;3000). Secondary hybridization was performed using fluorescently tagged antibodies: AF647 goat anti-rabbit (A2l24S, Invitrogen, l:l000), AFS94 goat anti-rabbit (All037, Life Technologies, l:l000), AFSSS goat anti-mouse (A28180, Invitrogen, l:l000) or AFS94 goat anti-mouse (All00S, Life Technologies, l:l000). Stained and post-fixed cells were imaged on a Nikon TI-E2 equipped with a Lasercombiner, a 60X SR P-Apochromat IR AC 60x 1.27 NA water immersion objective, a CSU-W1 SoRa spinning disk unit and an 0rca Fusion CM0S camera in spinning disk mode, operated using NIS Elements 5.2.02 (Nikon). Per condition (WT / ΔP0I), at least 5 z-stacks covering a R0I size of 26l.46×26l.46×2l µm were acquired in the DAPI channel (405 nm excitation), GFP channel (488 nm excitation, degradation control), and immunofluorescence channels (561 or 640 excitation) with a pixel size of about 227 nm in xy and 500 nm in z.

### Image Analysis of mitotic exit degradation samples

Image analysis of 3D stacks of stained mitotic exit cells was performed with a custom-written Python script. In brief, after a mild gaussian blur, the DAPI channel was converted to a 3D binary mask of nuclei used for 3D segmentation (method = triangle). Small objects and cropped nuclei at the image borders were removed automatically, and further quality control to remove poorly segmented, multinucleate or dead cells were removed manually using napari. Interactive viewing of the nuclei images and binary masks via napari was also used to classify cells as “mitosis” or “interphase” (representing all nuclei past anaphase). After classification, the nuclei mask and labels were used to extract fluorescent intensities of the endogenous P0I-GFP, as well as stained proteins in the unprocessed 488 nm, 561 nm and 647 nm (if applicable). Image background from regions devoid of cells was subtracted from mean nuclear pixel intensities in every image channel.

### Spot-bleach assay and analysis

Cells for spot-bleach measurements were seeded at a density of 2.Sxl0^S^ cells/ml into Ibidi glass bottom µ-Slide channels (80607, Ibidi) and grown for 16-24 hours. 0ne hour before imaging, DMEM was replaced by C0_2_-independent imaging medium (as above) containing S0-l00 nM S-SiR-Hoechst. FRAP experiments were performed on a LSM880 laser-scanning microscope with an inverted Axio 0bserver controlled by ZEN 2.1 Black software (Version l4.0.9.20l, Zeiss), equipped with an in-house constructed incubation chamber for temperature control set to 37°C and using a C-Apochromat 40x/1.2 W Korr UV-Vis-IR water-immersion objective (42l767-997l-7ll, Zeiss). Cells were screened at low-resolution live imaging in the SiR-Hoechst channel and image acquisition was started once a cell undergoing anaphase onset was identified. At S, l0, lS, 20 and 30 minutes after anaphase onset, an image of the dividing cell in the GFP (488 nm emission) and DNA (SiR-Hoechst, 633 nm emission) was acquired and used to place and initiate a 30 second continuous illumination with a diffraction limited focused laser beam (488 nm, ∼l.S µW laser power, corresponding to 0.1% Argon laser power). This resulted in a clear depletion of the chromatin-bound (m)EGFP-tagged protein pool and minor bleaching of the overall cellular pool that readily replaced the bleached soluble fraction at the measured spot. Measurement timepoints were distributed between the two daughter cells to further minimize light exposure of a single cell. During the 30 seconds illumination, emitted fluorescence was continuously measured using the GaAsP detector in photon counting mode. The mean of the first (prebleach) and last (postbleach) 500 milliseconds of the fluorescence depletion trace was used to calculate the chromatin-bound fraction for each measurement based on the following formula:

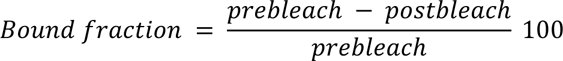

In addition to the measurements shortly after mitosis, chromatin-bound fractions of each P0I were measured in asynchronous interphase cells. Measured bound fractions were calibrated using exogenously H2B-EGFP (low expression level, positive control representing ∼100% chromatin bound fraction) and freely diffusing mEGFP (unbound control, representing 0% chromatin bound fraction) expressed in a HK WT cell background and measured in asynchronous interphase cell nuclei. The average calibrated chromatin-bound fractions of 10 spot-bleach measurements per protein per timepoint was interpolated (the asynchronous interphase measurements were set to 300 minutes after anaphase onset for this purpose) and used to calculate the average number of chromatin-bound P0Is at each timepoint during mitotic exit using the FCS-calibrated protein number information from Fig. 1EF, Suppl. Fig. 1P.

### Cell synchronization and immunofluorescence for chromatin tracing

To prepare HK cells expressing AID-EGFP-tagged Cohesin-STAG2 as well as HK WT cells for chromatin tracing in interphase, 120 µL of asynchronous AID-tagged and WT cells were seeded at a l:l ratio and a total density of Sxl0^S^ cells/ml into PBS-washed channels of Ibidi µ-Slide glass bottom slides (80607, Ibidi) and cultured for 20 hours at 37°C, S% C0_2_ in DMEM supplemented with 40 μM BrdU/BrdC (ratio 3:1, BrdU: B5002, Sigma-Aldrich, BrdC: sc-284SSS, Santa Cruz Biotech). Degradation of EGFP-AID-STAG2 was induced by the addition of 500 µM Inole-3-acetic acid (IAA, ISl48, Sigma-Aldrich) for 2 hours at 37°C, S% C0_2_ in DMEM. Cells were then fixed using 2.4% PFA (lS7l0, EMS) in PBS for lS minutes, followed by quenching of PFA with 100 mM NH_4_Cl in PBS (5 minutes) and washing with PBS. To prepare cells in early G1, HK WT and STAG2-AID cells were grown for 20 hours in a T-l7S flask (3S3ll2, Corning) in the presence of 40 μM BrdU/BrdC (ratio 3:1) to reach a confluency of around 80% suitable for mitotic shake off. Nocodazole-arrest, mitotic shake-off and resuspension of mitotic cells was performed as described above. Enriched mitotic HK WT and STAG2-AID cells (Wutz et al., 2020) were diluted to 2.Sxl0^6^ cells/ml, mixed l:l and 3S μL of this cell suspension was seeded into Ibidi µ-Slide glass bottom slides (80607, Ibidi) pre-coated with poly-L-lysine and incubated for lS minutes at 37°C, S% C0_2_ to allow cells to attach. Degradation of STAG2 was induced upon addition of 500 µM IAA in the presence of Nocodazole, ensuring near-complete degradation within 45 minutes. Release into mitotic exit was triggered by Nocodazole washout using DMEM containing 500 µM IAA. Cells were fixed 80 minutes after release. Live imaging of cells at this point showed that they are on average about 45 minutes past anaphase. After fixation, early G1 and asynchronous interphase cells were permeabilized for lS minutes using 0.2S% TritonX-100 (T8787, Sigma-Aldrich) in PBS and 0.l µm Tetraspec beads were added to the Ibidi channels (l:l00 dilution from stock, T7279, Thermo Fisher) to be used as fiducials for drift correction. After blocking with 2% BSA in 0.05% TritonX-100 at RT for at least 30 minutes, primary labelling of STAG2 was performed overnight at 4°C in a humidified chamber (with rabbit-anti STAG2, Glycine Elution, 1:200, Sumara et al., 2000), followed by hybridization with an AF488-labelled secondary antibody (goat-anti-rabbit AF488, A-ll034, Molecular Probes).

### Non-denaturing FISH (RASER-FISH)

Non-denaturing FISH (RASER-FISH) as well as FISH library design and amplification was performed as described previously (Beckwith et al., 2023 *Preprint* & Beckwith, Brunner et al., in preparation). In brief, cells were incubated with 0.5 ng/µl DAPI in PBS at RT for lS minutes to sensitize DNA for UV-induced single-strand nicking of the replicated strand containing BrdU/C. Subsequently, the cells were exposed (without Ibidi lid) to 254 nm UV light for lS min (Stratalinker 2400 fitted with lSW 2S4 nm bulbs-part no GlST8). The nicked strand of DNA was then digested using Exonuclease (lU/ul, M0206, NEB) in NEB buffer 1 at 37 °C for lS min in a humidified chamber. Cells were post-fixed using S mM Bis(NHS)PEGS (803S37, Sigma-Aldrich) in PBS for 30 minutes at RT to preserve cell fixation during primary FISH library hybridization at 37°C. Hybridization of primary FISH probe libraries targeting 1.2 Mb regions (Chrl4 S0.92-S2.l0 Mb, ChrS l49.S0-lS0.70 Mb, Chr2 l9l.ll-l92.3l Mb) with 12 kb genomic resolution (one trace-spot = tiled set of ∼lS0 FISH probes with common docking handle) was performed by incubation with hybridization buffer (50% formamide (FA, AM9342, Thermo Fisher), 10% (w/v) dextran sulfate (D8906, Sigma-Aldrich) in 2xSSC (AM9763, Thermo Fisher) containing the FISH probe libraries at a final concentration of 100-200 ng/µL DNA per library for 1-2 nights at 37°C in a humidified chamber. After primary hybridization, channels were rinsed 3 times with 50% FA in 2xSSC, washed again twice with 50% FA in 2xSSC for 5 min at RT and finally washed with 2xSSC containing 0.2% Tween. RNA-DNA hybrids were removed by incubating cells with 0.05 U/µL RNAse H (M0297S, NEB) for 20 min at 37 °C in RNAse H buffer (NEB). To image and segment whole 1.2 Mb tracing loci, secondary FISH probes serving as bridges between all primary probes of a whole 1.2 Mb locus and a common imager strand were applied at a concentration of l00 nm in secondary hybridization buffer (20% Ethylene Carbonate (EC, E26258, Sigma-Aldrich), 2xSSC) for 20 minutes at RT rocking. Secondary probes were then washed with 30% FA in 2XSSC at RT (3 washes, 5 minutes each) and 2 additional washes with 2xSSC. Prior to imaging, DNA was stained with 0.5 ng/µl DAPI in PBS for 5 minutes at RT.

### Chromatin Tracing using LoopTrace

3D DNA trace acquisition using a custom-built automated fluidics setup was performed as described in Beckwith et al., 2023 (*Preprint*) and in https://git.embl.de/grp-ellenberg/tracebot. In brief, 12-mer imager strands with 3’ or S’-azide functionality (Metabion) complementary to the docking handles employed by the primary FISH probe library, as well as the bridged regional barcode probes added during secondary hybridization, were fluorescently labelled with Cy3B-alkyne (AAT Bioquest) or Atto643-alkyne (Attotec) using click chemistry (ClickTech 0ligo Link Kit, Baseclick GmbH) according to the manufacturer’s instructions to enable dual-color tracing. Fluorescently labelled 12-mer imagers were diluted to a final concentration of 20 nM in 5% EC 2X SSC in a 96 well plate and placed on the stage of a custom-built automated fluidics setup based on a GRBL controlled CNC stage (Beckwith et al., 2023). Furthermore, a 3-well deep plate containing washing buffer (10 % FA, 2X SSC) and stripping buffer (30% FA, 2XSSC) covered with parafilm, as well as a 24-well plate containing imaging buffer (0.2X Glucose 0xidase (G7l4l, Sigma-Aldrich), l.S mM TR0L0X (2388l3, Sigma-Aldrich), 10% Glucose, 50 mM Tris, 2X SSC pH 8.0) were placed on the stage of the automated fluidics setup. A syringe needle mounted in place of the CNC drill head was connected to the sample and a CPP1 peristaltic micropump (Jobst Technologies, Freiburg, Germany, flow rate of 1 mL/min at maximal speed) using 1 mm i.d. PEEK and silicone tubing (VWR), allowing to pull liquids out of the well plates and through the sample channel in an automated manner. Imaging was performed on a Nikon TI-E2 microscope equipped with a Lasercombiner, a l00X l.3S NA silicon oil immersion objective, a CSU-W1 SoRa spinning disk unit and an 0rca Fusion CM0S camera in spinning disk mode, operated using NIS Elements 5.2.02 (Nikon) in combination with custom-made Python software for synchronization with automated liquid handling. Prior to sequential imaging, a 3D stack of DAPI-stained nuclei (405 nm excitation), STAG2-EGFP fluorescence (488 nm excitation) and the fiducial beads (561 or 640 excitation) was acquired as a reference stack for cell classification with a pixel size of 130 nm in xy and 300 nm in z at a total size of l49.76xl49.76 µm in xy and covering a z-range of l4.l (interphase) - 18.3 µm (early G1). Subsequently, imager strands were sequentially hybridized for ∼ 2 minutes at 20 nM concentration in 5% EC 2X SSC, washed for 1 minute with washing buffer, imaged after addition of GL0X-based imaging buffer as a 3D stack, stripped for ∼2 minutes using stripping buffer and washed again for 1 minute. 3D stacks acquired during sequential imaging had equal pixel sizes and z-range as before, but were acquired only in the 561 nm or 640 nm channels (100% laser power, 100 msec exposure time, triggered acquisition mode), to image fiducial beads and Cy3B or Atto643-labelled imagers, respectively.

### Analysis of LoopTrace data

Processing of acquired tracing data was performed as described in Beckwith et al., 2023 (*Preprint*) with code available under https://git.embl.de/grp-ellenberg/looptrace. In brief, nd2 image files were converted to 0ME-ZARR format. Images were drift-corrected based on cross-correlation and sub-pixel drift was corrected by fitting the fiducial bead signal to a 3D gaussian function and subsequent correction for calculated sub-pixel drift. Images were deconvolved using the experimental PSF extracted from fiducial beads. Identification of tracing regions was performed based on regional barcodes using an intensity threshold. Detected spot masks were then used to extract regions of interest for 3D-superlocalisation of individual trace-spots by fitting with a 3D gaussian. Finally, extracted traces were corrected for chromatic aberration between the 561 and 642 image channels by affine transformation obtained by least squares fitting of the centroid of fiducial beads imaged in both channels, and traces were assigned to nuclei classified as “interphase”, “early Gl” or “mitosis”.

The resulting interphase and early G1 DNA traces were grouped into “WT” or “ΔSTAG2” based on their AF488 intensity and the subsequent analysis was performed as described in Beckwith et al., 2023 (*Preprint*). In brief, all fits were quality-controlled for their signal to background ratio, standard deviation of the fit and fit center distance to the regional barcode signal. Traces containing less than 20 high-quality fitted positions were removed from further analysis. Median pairwise distances were calculated for all 3D coordinates within a single trace and used to display either pairwise-distance maps or contact maps by calculating the frequency of contacts below a certain 3D distance (set to 120 nm). Difference matrices were achieved by subtraction of “dSTAG2” from “WT” pairwise distances. Scaling plots were generated from pairwise distance metrices as well, essentially plotting all measured 3D distances for every given genomic distance.

### Sample preparation for STED microscopy

To prepare genome-edited HK cells expressing endogenously EGFP-tagged Cohesin-STAG1/2 for STED microscopy in early Gl or Gl, cells were synchronized in mitosis and subsequently released into mitotic exit, pre-extracted, PFA-fixed and immuno-stained. Cell synchronization was performed by mitotic shake-off as described above. 35 µL of Nocodazole-arrested enriched mitotic cells were added at a density of l.2xl0^6^ cells/ml (for Gl) or 2.Sxl0^6^ cells/ml (for early G1) to pre-washed and poly-L-lysine coated channels of Ibidi µ-Slide glass bottom slides (80607, Ibidi) and incubated for lS minutes at 37°C, S% C0_2_ to allow cells to attach. Subsequently, 3 washes with fresh DMEM were performed to wash out Nocodazole, and cells were allowed to exit mitosis for 45 minutes (for early G1 stage) or 4h (for G1 stage) at 37°C, S% C0_2_. Pre-extraction was performed by washing cells once in PBS and then adding 0.25% Tergitol in 1X PBS for a total of 1 minute. Cells were then immediately fixed using 2.4% PFA in PBS for lS minutes, followed by quenching of PFA (lS7l0, EMS) with 100 mM NH_4_Cl in PBS and washing with PBS. Fixed cells were prepared for immuno-staining by an additional lS-minute permeabilization (standard IF protocol) in PBS with 0.25% Tergitol and subsequent blocking using blocking buffer (2% BSA in 0.05% Tergitol in PBS) for at least 30 minutes at RT. Incubation with the anti-GFP nanobody (FluoTag®-X4 anti-GFP conjugated to Abberior® Star 63SP, l:2S0 dilution N0304-Ab635P, NanoTag) and rabbit anti-CTCF antibody (Glycine-Elution, 1:3000, Wutz et. al 2020) was performed in blocking buffer at 4°C in a humidified chamber overnight. Secondary hybridization using AF594-conjugated goat-anti-rabbit antibody (l:l000, All037, Life Technologies) was performed for 1h at RT. Samples were post-fixed for lS minutes in 2.4% PFA in PBS, with subsequent quenching (100 mM NH_4_Cl in PBS) and PBS washing. Samples were imaged by STED super-resolution microscopy on the same day.

### STED microscopy

2D STED imaging was performed on a Leica Stellaris 8 STED Falcon FLIM microscope (Leica Microsystems) controlled by the Leica LAS X software (4.7.0.28176). Samples were imaged at RT using a HC PL AP0 86x/l.2 W motC0RR STED white water immersion objective. The microscope was equipped with the SuperK FIANIUM FIB-12 white light laser with laser pulse picker (440-790 nm, Leica Microsystems/NKT), 592 nm continuous wave (cw), 660 nm cw and 775 nm pulsed lasers (MPB Communications) and the HyD S, HyD X and HyD R detectors. Diffraction-limited as well as STED imaging of CTCF (AFS94) and STAGl/2-EGFP (Abberior Star 635P) was performed with excitation at S90 nm and 64S nm using the white light laser (diffraction-limited/confocal: 3% each, STED: 590 nm: 9%, 645 nm: 6%). Fluorescence was detected with two HyD X detectors using a 60l-6l9 nm and a 6SS-7S0 nm detection window, respectively. Imaging was performed in xy line sequential mode. The pinhole size was set to 1 airy unit and the pixel size set to l8.88xl8.88 nm in xy, resulting in images capturing a region of l9.3lxl9.3l µm with l024xl024 pixels. STED imaging was performed using a 2D depletion doughnut and 50% power with a 775 nm depletion laser for super-resolved imaging of CTCF-AFS94 and at l2% excitation power at 775nm for imaging of STAG1/2-Abberior Star 635P. STED images were acquired using 16 line accumulations with a scan-speed of 200 Hz resulting in a pixel dwell time of 3.85 µs. STED imaging was performed in FLIM mode and the images were post-processed using tau-STED enhancement with background suppression activated and tau-strength set to 0%. Crosstalk between fluorescent channels was quantified with the settings described above and found to be less than 5% (maybe ref to supplementary figure, if space allows).

### STED image analysis

Pre-processing of diffraction-limited and STED images was performed using Fiji using a custom-made script. Nuclear masks were created by segmenting the CTCF-AF594 diffraction-limited image after gaussian-blurring and used to crop out nuclei in all image channels. In addition, STED images were background subtracted using the rolling ball algorithm set to a radius of 50 pixels.

Co-localization analysis, as well as spot segmentation was performed using custom python scripts. For co-localization analysis, Pearson correlation coefficient of CTCF and STAG1/2 was computed based on cropped nuclei in the respective STED channel. Spot segmentation was performed by first coarsely segmenting spots inside the nuclear mask based on a common threshold (method: 0tsu) after applying a mild gaussian blur (sigma = l). Image noise resulting in excess tiny spots was filtered out through binary mask erosion and filtering, followed by binary dilation of correctly detected spots. Coarsely segmented spots often represent clusters of spot signals and were further segmented using a combination of local peak finding & watershed. The resulting masks for individual spots were used to extract average pixel intensities in the STED and confocal images. Assuming a z-depth of about 500 nm, the number of detected spots per μm^3^ was compared to protein number estimates derived by FCS-calibrated imaging of early Gl or Gl cells to estimate the overall labelling efficiency.

### STED image simulation

STED images were simulated by generating a desired number of randomly localized spots (single pixels) in an image representing 200 μm^2^ (or 100 μm^3^ assuming a z-depth of 500 nm), given the pixel size of 18.88 nm in the images acquired as described above. The randomly distributed spots were gaussian blurred (sigma = 2.6) and their pixel intensity was enhanced 6-fold to be distinguishable above a random background. Simulated images were analyzed for co-localization or segmented and analyzed to read out their average spot intensities as described above.

