## Supplementary Information for "Quantitative imaging of loop extruders rebuilding interphase genome architecture after mitosis"

### This PDF file includes:

Supplementary Figures

Supplementary Tables 1&2

### Other Supplementary Materials for this manuscript include the following:

- List of antibodies
- List of cell lines
- List of gRNAs

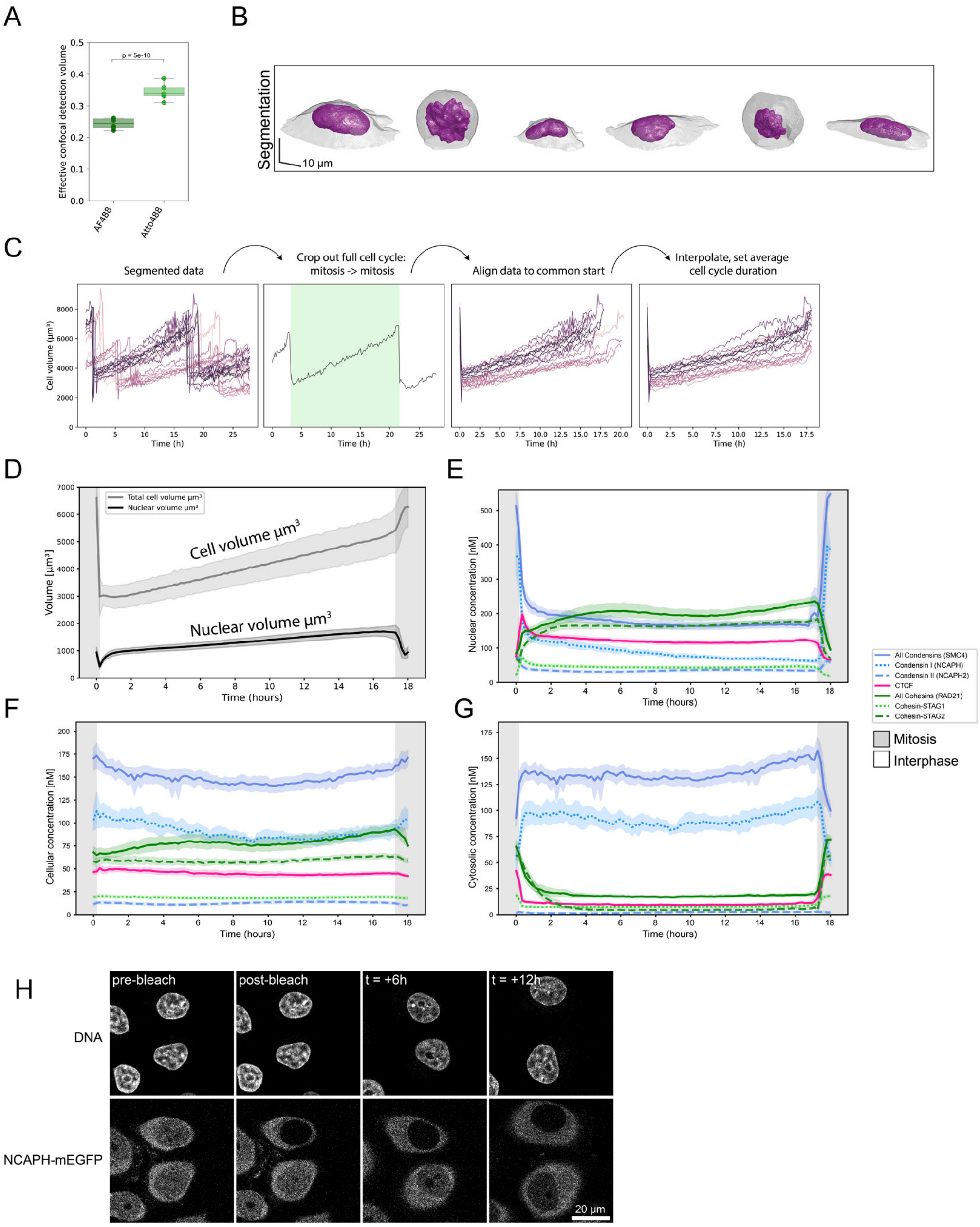


**Supplementary Figure 1, page 1**

**A)** Effective confocal detection volume ($V_{eff}$) determined using AF488-NHS dye (Politi et al., 2018) and Atto488 (this study) at low concentrations (10 nM). 5 30-second-long FCS measurements are performed for $V_{eff}$ determination. Autocorrelation analysis of these measurements and fitting of the diffusion time parameter $\tau_D$ is performed to calculate a 3D-gaussian volume.

**B)** Exemplary segmentation of a full cell cycle track. Segmentations correspond to the top right cell in Fig. 1C.

**C)** Illustration of full cell cycle data processing based on cell volume information.

**D)** Average cell and nuclear volume for all single cell trajectories combined. Error bands represent standard deviation.

**E-G)** Mean nuclear (E), cellular (F) and cytosolic (G) concentrations of HeLa Kyoto homozygous knock-in cell lines.

**H)** The nuclear NCAPH pool of cells endogenously expressing NCAPH-EGFP was photobleached and monitored for up to 12 hours. Full bleaching could even be achieved with bleach ROIs that do not target the entire nuclear volume, indicating fast and freely moving protein. The bleached nuclear pool was not recovered by unbleached cytosolic pool throughout the measurement period.

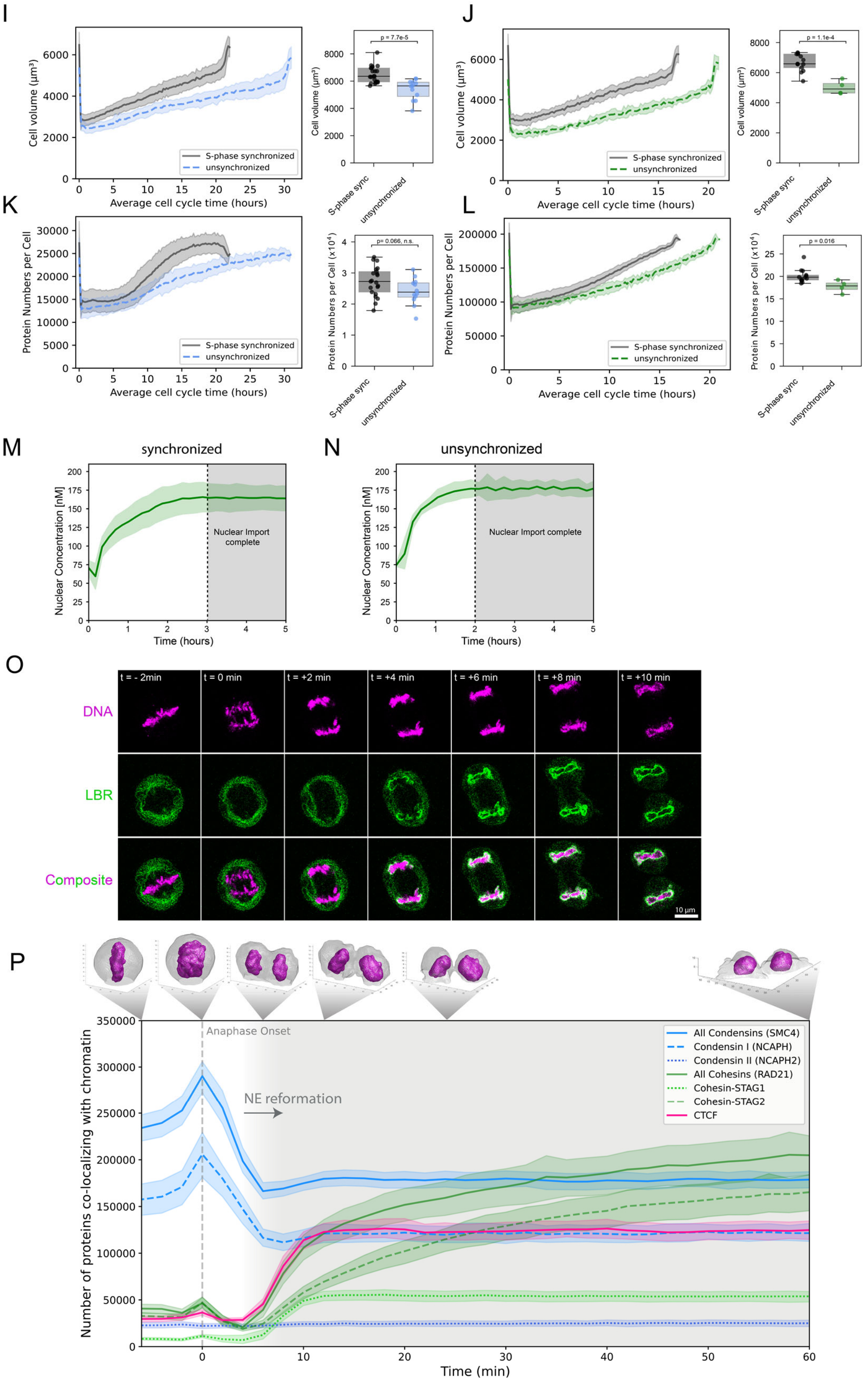


**Supplementary Figure 1, page 2**

**I-L)** Comparison of cell volume and absolute protein numbers in S-phase synchronized versus asynchronous cells. Bar plots compare cell volume or cellular protein content in the first mitosis after release from S-phase arrest. S-phase arrest resulted in notably increased cell size and protein abundance, influencing protein

abundance, production and duration of the next cell cycle. See L&M for the influence of synchronization on Cohesin-STAG2 protein import.
**M-N)** Comparison of Cohesin-STAG2 protein import kinetics in S-phase synchronized versus asynchronous cells. In contrast to S-phase synchronized cells, we found that WT cells required only about 2 hours for full Cohesin-STAG2 import into the newly formed nuclei.
**O)** Live cell imaging of the nuclear envelope marker LBR-GFP (Ellenberg et al., 1997) to assess the timing of nuclear envelope reformation after mitosis. While LBR-GFP chromatin (stained via 5-SiR-Hoechst) accumulated as early as +4 minutes after anaphase onset (AO), chromatin was almost fully engulfed at +6 minutes past AO and fully covered at +10 minutes past AO.
**P)** Absolute protein numbers co-localizing with chromatin/the two daughter nuclei displayed for genome-edited HK cells with homozygously (m)EGFP-tagged proteins (SMC4: n = 21 cells, NCAPH: n = 14 cells, NCAPH2: n = 19 cells, CTCF: n = 15 cells, RAD21: n = 18 cells, STAG1: n = 25 cells, STAG2: n = 11 cells). Reformation and full establishment of the nuclear envelope as determined by Lamin B receptor (Suppl. Fig. 1N) is indicated through grey background. Error bands represent 95% confidence interval.

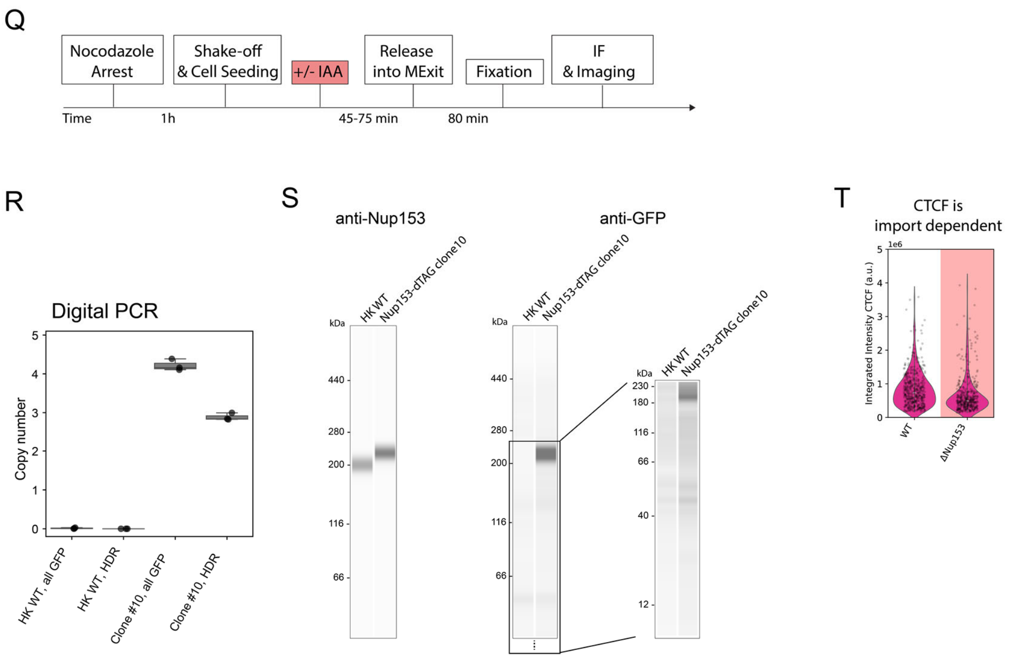


**Figure 1 supplement, page 3.**

**Q)** Experiment scheme for mitotic synchronization of genome-edited HK cells with homozygously mEGFP-FKBP12$^{F36V}$ tagged Nup153, followed by targeted protein degradation of Nup153, release into mitotic exit and timed fixation during early G1. Cells are immune-stained for RAD21 or CTCF and for diffraction-limited imaging.

**R-S)** Validation data for the generation of the homozygous Nup153-mEGFP-FKBP12$^{F36V}$ knock-in line, clone #10.

**R)** Digital PCR was performed to assess the copy number of GFP inserted into the HK genome, and to assess the number of successful homology directed repair (HDR) events at the Nup153 endogenous gene locus. While one copy of mEGFP-FKBP12$^{F36V}$ was inserted elsewhere in the genome, this copy is not expressed (T).

**S)** Simple Western analysis of HK WT cells and the Nup153-mEGFP-FKBP12$^{F36V}$ #C10 cell line created and used within this study. The anti-GFP western blot shows no expression of free GFP. The anti-Nup153 western blot shows a clear shift of the Nup153 band, indicating successful and homozygous gene editing.

**T)** Average CTCF fluorescence intensity upon immunostaining in early G1 cells in WT condition or after mitotic depletion of Nup153. ΔNup153 cells show a 25-40% reduction in average CTCF fluorescent intensity after 45 min release time. Changes above/below 20% are considered a significant change.

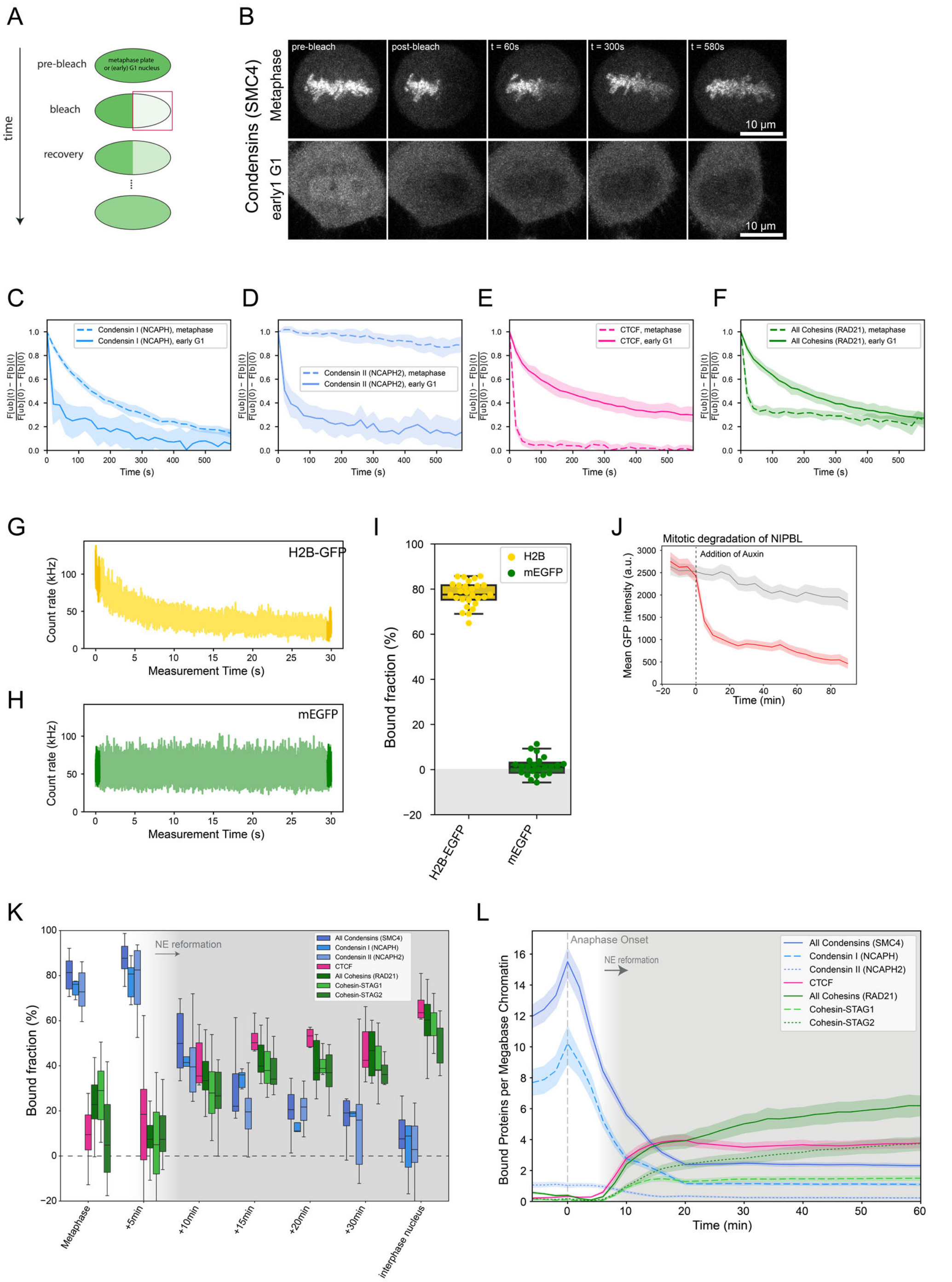


**Supplementary Figure 2.**

**A)** Scheme of FRAP experiments. Half of the metaphase plate/nucleus was bleached and fluorescent recovery was monitored in bleached and unbleached regions.

**B)** Exemplary FRAP data of SMC4-mEGFP in metaphase and early G1 cells. While 1 bleach step (150 repetitions 100% laser power) is performed in metaphase cells, 3 bleach steps (50 repetitions, 100% laser power each) are performed in early G1 cells to bleach the entire soluble pool and allow for the determination of the total chromatin-bound fraction.

**C-F)** Metaphase and early G1 FRAP measurements of Condensin I (NCAPH-mEGFP, C), Condensin II (NCAPH2-mEGFP, D), CTCF-EGFP (E) and Cohesin (RAD21-EGFP, F).

**G)** Spot-bleach measurement of HK WT cells exogenously expressing low concentration of stably chromatin-bound H2B-EGFP. H2B-EGFP chromatin bound fraction was used to calibrate spot-bleach measurements.
**H)** Spot-bleach measurement of HK WT cells exogenously expressing low concentration of monomeric EGFP. mEGFP chromatin bound fraction was used to calibrate spot-bleach measurements.
**I)** Chromatin-bound fractions of H2B-EGFP (used as calibration reference for 100% chromatin-bound) and mEGFP (used as calibration reference for 0% chromatin-bound pool). Chromatin-bound fractions of all other proteins of interest were scaled accordingly.
**J)** Mean cellular fluorescence intensity of endogenous NIPBL tagged with AID-EGFP in WT condition or upon addition of auxin measured in Nocodazole-arrested mitotic cells. Depletion of the protein pool happened within 20 minutes. Low signal/background ratio required high light-doses leading to bleaching of NIPBL-EGFP and autofluorescence.
**K)** The fraction of chromatin-bound Condensin and Cohesin isoforms as well as CTCF determined using the spot-bleach assay at different timepoints during mitotic exit. Every bar plot represents at least 10 individual datapoints measured in 10 separate cells.
**L)** Absolute number of proteins bound to chromatin were determined by multiplication of chromatin bound fractions shown in (K) with absolute protein numbers co-localizing with chromatin as determined in Fig. 1E&F & Suppl. Fig. 1P and displayed as per-megabase-count assuming an equal distribution of the proteins on the entire 7.9 Mb HeLa genome (Landry et al., 2013). Grey background indicates reformation of nuclear envelope. Error bands represent 95% confidence interval.

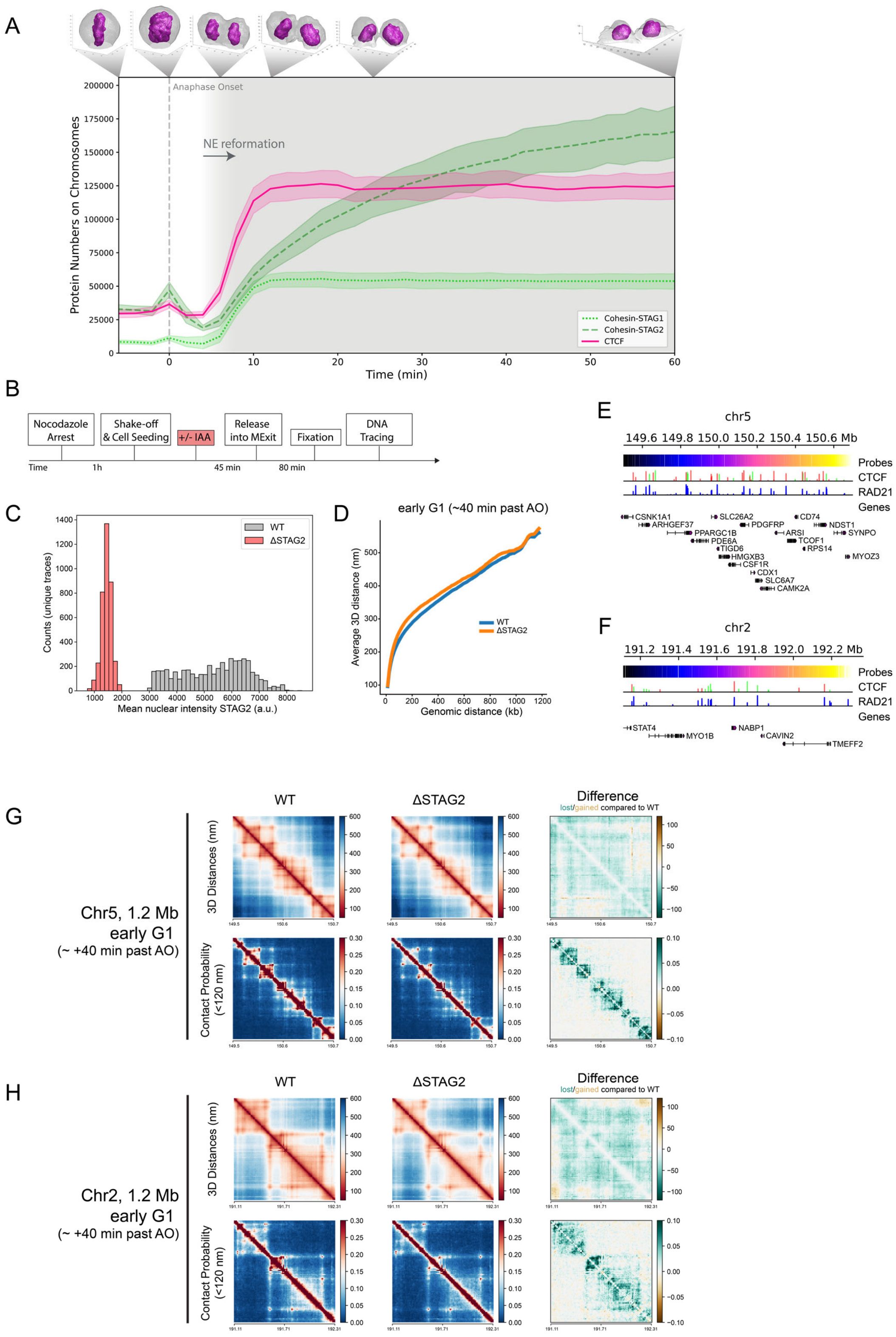


**Supplementary Figure 3.**

**A)** FCS-calibrated protein numbers co-localizing with chromatin are displayed for genome-edited HK cells with homozygously EGFP-tagged Cohesin-STAG1 (n = 25 cells), Cohesin-STAG2 (n = 11 cells) and CTCF (n = 15 cells). Error bands represent 95% confidence interval.

**B)** Experimental scheme for mitotic degradation of Cohesin-STAG2 using genome-edited HK cells with homozygously AID-EGFP tagged Cohesin-STAG2, followed by release into mitotic exit and chromatin tracing using LoopTrace (Beckwith et al., 2023 *Preprint*).
**C)** Mean nuclear intensity of immune-stained STAG2 in WT condition or after depletion of endogenously tagged STAG2. Nuclei and the corresponding traces could be clearly classified into WT or ΔSTAG2.
**D)** Scaling plot of Chr14 1.2 megabase region sampled at 12 kb resolution in WT or ΔSTAG2 cells. Traces from ΔSTAG2 are slightly less compact compared to WT.
**E-F)** Overview of the traced 1.2 megabase locus on chromosome 5 (E) and chromosome 2 (F) with genes as well as ChIP-seq binding sites for RAD21 and CTCF (from the ENCODE portal (Sloan et al., 2016), https://www.encodeproject.org/) with the following identifiers: ENCFF239FBO (RAD21), ENCFF111RWV (CTCF); CTCF directionality annotations from Rao et al., 2014).
**G-H)** Distance and contact matrices of a 1.2 megabase region on chromosome 5 (G) and chromosome 2 (H) traced at a genomic resolution of 12 kb in early G1 cells with and without Cohesin-STAG2. Differences between WT and ΔSTAG2 are highlighted for distance and contact probability maps.

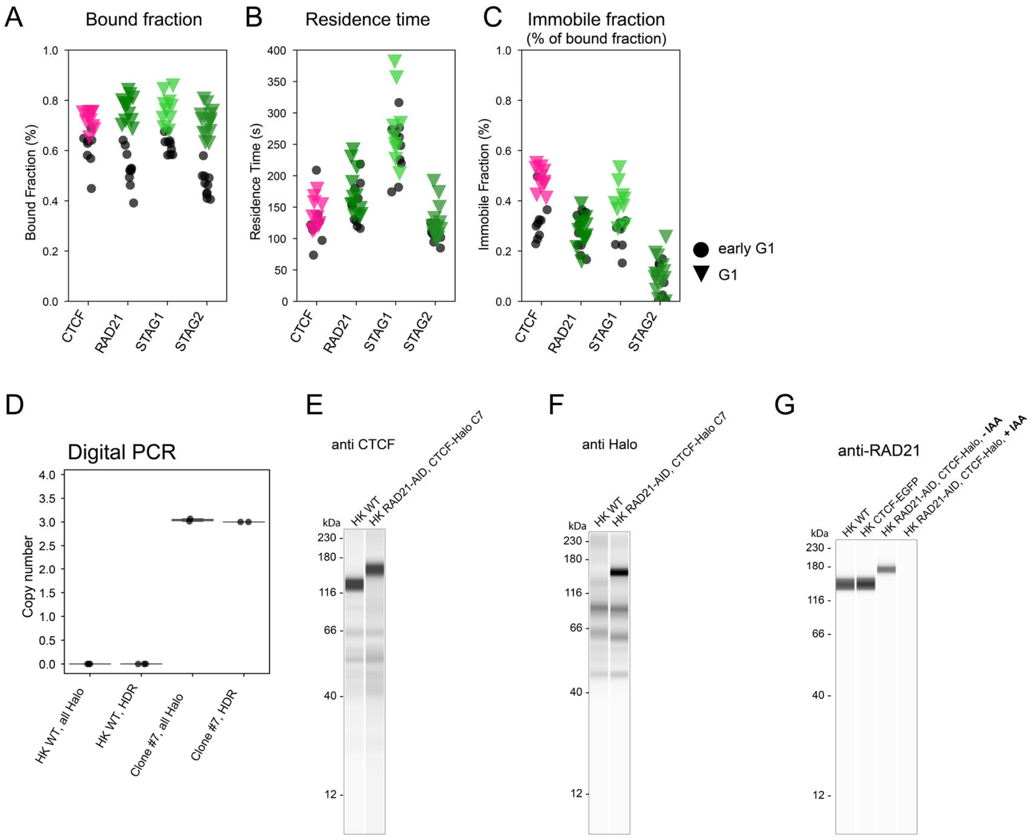


**Supplementary Figure 4.**

**A-C)** Chromatin-association metrics derived from FRAP measurements in HK cells in which CTCF, RAD21, STAG1 and STAG2 were endogenously tagged with EGFP. Comparison between early G1 and later G1 measurement timepoint.

**A)** Chromatin bound fractions were calculated based on the remaining fluorescent intensity in the unbleached region after bleaching. After the 3$^{rd}$ bleach iteration the entire soluble nuclear protein pool was bleached. Differences between early G1 and G1 were significant for all proteins tested given a significance level of p = 5% (Kolmogorov-Smirnov test, CTCF: p = 0.0037, RAD21: p = $3.09 \times 10^{-6}$, STAG1: p = 0.0002, STAG2: p = $1.75 \times 10^{-6}$).

**B)** Chromatin residence times were derived by fitting FRAP recovery with a single exponential function with an immobile fraction component. Differences between early G1 and G1 were non-significant for all proteins tested given a significance level of p = 5% (Kolmogorov-Smirnov test, CTCF: p = 0.39, RAD21: p = 0.28, STAG1: p = 0.54, STAG2: p = 0.22).

**C)** Immobile fractions were derived by fitting FRAP recovery with a single exponential function with an immobile fraction component. Differences between early G1 and G1 were significant for CTCF and STAG1 given a significance level of p = 5% (Kolmogorov-Smirnov test, CTCF: p = 0.0004, RAD21: p = 0.5577, STAG1: p = 0.0037, STAG2: p = 0.1497.)

**D-F)** Validation data for the correct tagging of CTCF in the HK RAD21-EGFP-AID CTCF-Halo-3xALFA (#C7) cell line generated in this study.

**D)** The copy number of Halo-3xALFA tags integrated at the target locus (HDR assay) and within the whole recipient genome (all-Halo assay) was determined in HK WT and edited (clone #7) cell lines by digital PCR. The complete tagging of all 3 endogenous CTCF copies was confirmed by PCR-amplification of the target locus and sequencing analysis. Digital PCR results indicate that no extra off-target copies of the tag are present at the genome of edited cells.

**E-F)** Simple western analysis of protein extracts from HK WT cells and the RAD21-EGFP-AID CTCF-Halo-3xALFA #C7 line created and used in this study. For each condition, 3 µL of total protein lysate at 0,4 µg/µL was loaded into the assay's microplate.

**E)** Immunolabeling of CTCF shows a clear shift of the CTCF band to higher molecular weight in the edited cell line, indicating successful and homozygous gene tagging. Anti-CTCF antibody (07-729, EMD Millipore) was used at 1:40 dilution.
**F)** Immunolabeling of Halo shows correct tagging of a protein of the expected MW for CTCF-Halo-3xALFA, and no expression of free Halo tag. Anti-Halo Antibody (G9211, Promega) was used at 1:50 dilution.
**G)** Simple western analysis of protein extracts from HK WT cells, HK CTCF-EGFP cells and the RAD21-EGFP-AID CTCF-Halo-3xALFA #C7 line created and used in this study, the later grown in the absence (-IAA) and presence (+IAA) of auxin in the last 3 hours of culture. For each condition, 3 µL of total protein lysate at 0,4 µg/µL was loaded into the assay's microplate. Anti-RAD21 antibody (05-908, Sigma-Aldrich) was used at 1:50 dilution. Complete depletion of RAD21 in the genome-edited cell line was achieved by the addition of auxin. In the absence of auxin, this cell line showed a reduced expression of RAD21 compared to HK WT or the HK CTCF-EGFP cell line due to leaky degradation of RAD21. Nonetheless, the effect of the loss of the remaining RAD21 in the double-knock-in cell line still led to a more dynamic interaction of CTCF with chromatin (assessed by FRAP, data not shown).

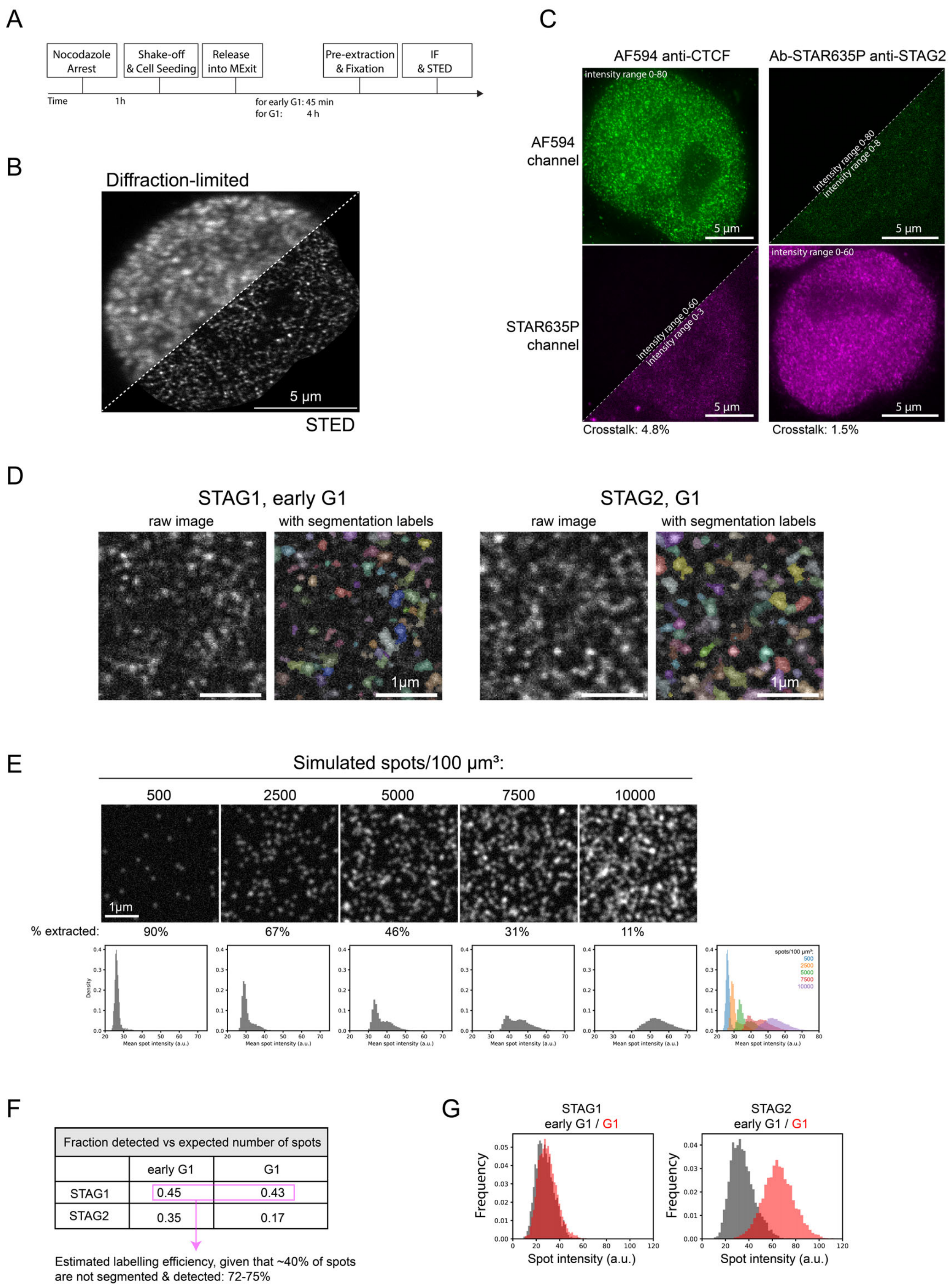


**Supplementary Figure 5, page 1**

**A)** Experimental scheme for mitotic synchronization of genome-edited HK cells with homozygously EGFP tagged Cohesin-STAG1/2, followed by release into mitotic exit, fixation during early G1 or G1, immunostaining for CTCF and STAG1/2-EGFP and subsequent STED super-resolution imaging.

**B)** Exemplary G1 cell expressing STAG1-EGFP, with Cohesin-STAG1 stained via a GFP nanobody. Imaged in confocal or STED mode, respectively.

**C)** Cross-correlation control of STED image channels. Cells in which either CTCF-AF594 or STAG1/2-EGFP-Abberior-STAR-635P were labelled. Cross-talk between imaging channels was assessed and found to be minimal (<5%).

**D)** Illustration of the STED image spot segmentation.

**E)** Benchmarking of spot-counting via the segmentation workflow (F). At higher spot-densities the true spot-counts are underestimated. Our STED imaging data appears to have very similar spot densities as the simulated images with 2500-5000 spots/µm$^3$ (for STAG1 and STAG2 in early G1). Given that only 67-46% of all spots are segmented and counted in these simulated images, we are likely also undercounting protein numbers in our acquired STED data by approximately 40%.
**F)** The number of segmented (= detected) spots per cubic micrometer (assuming a typical z-depth of 500 nm for STED, averaged for all three replicates) compared to the expected number of chromatin-bound STAG1/2 proteins per cubic micrometer as measured by FCS-calibrated imaging and FRAP (Tables 1&2). Given that the segmentation pipeline underestimates the number of proteins (H) by about 40% (compare density to images in Fig 5. A) the labelling efficiency can be estimated to about 72-75%. Note the 2-fold decrease in detected proteins spots for STAG2 from early to later G1.
**G)** Mean intensity of segmented STAG1/2 spots in confocal images of replicate 3. Same results are observed in replicate 1 and 2. Formal significance tests are meaningless due to large sample size.

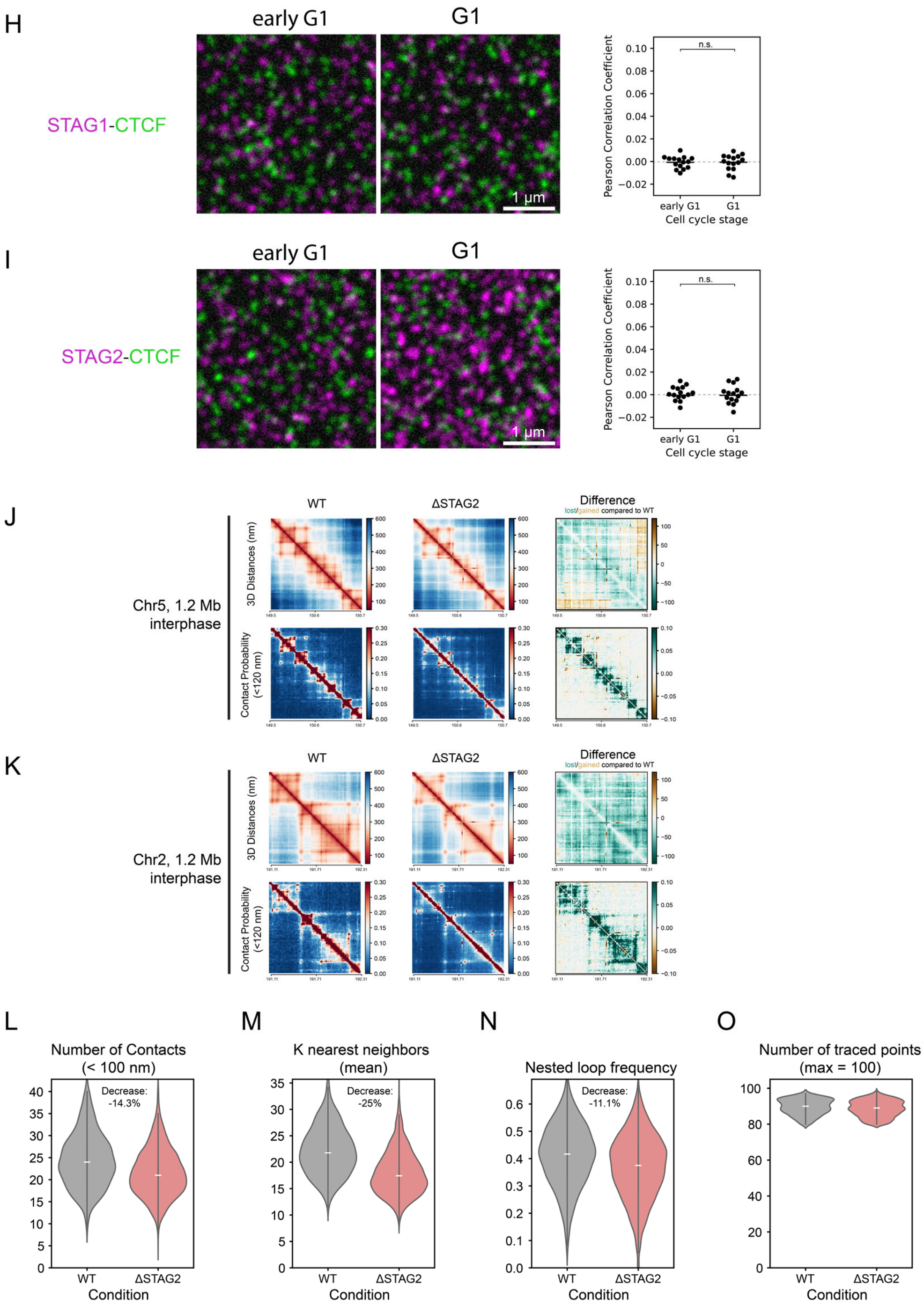


**Supplementary Figure 5, page 2**

**H)** Image simulation based on detected spot densities in acquired STED images for STAG1 and CTCF, multiplied by 1.67 to account for the fact that the spot-segmentation workflow underestimates STED spot densities around 40%.

**I)** Image simulation based on detected spot densities in acquired STED images for STAG2 and CTCF, multiplied by 1.67 to account for the fact that the spot-segmentation workflow underestimates STED spot densities around 40%.

**J-K)** Distance and contact matrices of a 1.2 megabase region on chromosome 5 (J) and chromosome 2 (K) traced at a genomic resolution of 12 kb in interphase cells with and without Cohesin-STAG2. Differences between WT and ΔSTAG2 are highlighted for distance and contact probability maps. WT data from 1, ΔSTAG2 data from 2 independent technical replicates (>400 traces, respectively).

**L-O)** Trace metric analysis from single chromatin traces of 1.2 Mb regions on chromosome 14, chromosome 5 and chromosome 2 (data from all 3 loci was combined to show one average plot per trace metric). Number of contacts below a physical distance of 100 nm (L), the mean number of k nearest neighbors (M) and the frequency of nested loops (requirement: 2 base loops have a common anchor point and thereby form a bigger stacked loop, N) are shown for >1000 individual traces per condition that are at least 80% or more complete (O).

Supplementary Table 1. Bound fraction comparison FRAP vs spot-bleach assay, **early G1**

| Early G1 | | |
|---|---|---|
| **Protein of interest** | **% bound (spot-bleach, average)** | **% bound (FRAP)** |
| **SMC4** | 0.21 | 0.19 |
| **NCAPH** | 0.15 | 0.11 |
| **NCAPH2** | 0.22 | Optimal parameters for fitting not found |
| **CTCF** | 0.5 | 0.62 |
| **RAD21** | 0.41 | 0.53 |
| **STAG1** | 0.37 | 0.62 |
| **STAG2** | 0.37 | 0.47 |

Supplementary Table 2. Bound fraction comparison FRAP vs spot-bleach assay, **G1/interphase**

| G1/Interphase | | |
|---|---|---|
| **Protein of interest** | **% bound (spot-bleach, average)** | **% bound (FRAP)** |
| **CTCF** | 0.72 | 0.72 |
| **RAD21** | 0.59 | 0.77 |
| **STAG1** | 0.59 | 0.76 |
| **STAG2** | 0.5 | 0.71 |
